# Disease modification upon brief exposure to tofacitinib during chronic epilepsy

**DOI:** 10.1101/2023.08.07.552299

**Authors:** Olivia R. Hoffman, Jennifer L. Koehler, Jose Ezekiel Clemente Espina, Anna M. Patterson, Emily S. Gohar, Emanuel Coleman, Barry A. Schoenike, Claudia Espinosa-Garcia, Felipe Paredes, Nicholas H. Varvel, Raymond J. Dingledine, Jamie L. Maguire, Avtar S. Roopra

## Abstract

All current drug treatments for epilepsy, a neurological disorder affecting over 50 million people(*1, 2*) merely treat symptoms, and a third of patients do not respond to medication. There are no disease modifying treatments that may be administered briefly to patients to enduringly eliminate spontaneous seizures and reverse cognitive deficits(*3, 4*). Applying network approaches to rodent models and human temporal lobectomy samples at both whole tissue and single-nuclei resolutions, we observe the well-characterized pattern of rapid induction and subsequent quenching exhibited of the JAK/STAT pathway within days of epileptogenic insult. This is followed by a resurgent activation weeks to months later with the onset of spontaneous seizures. Targeting the first wave of activation after epileptic insult does not prevent disease. However, brief inhibition of the second wave with CP690550 (Tofacitinib) (*5, 6*) enduringly suppresses seizures, rescues deficits in spatial memory, and alleviates epilepsy-associated histopathological alterations. Seizure suppression lasts for at least 2 months after the final dose. Using discovery-based transcriptomic analysis across models of epilepsy and validation of putative mechanisms with human data, we demonstrate a powerful approach to identifying disease modifying targets; this may be useful for other neurological disorders. With this approach, we find that reignition of inflammatory JAK/STAT3 signaling in chronic epilepsy opens a window for disease modification with the FDA-approved, orally available drug CP690550.

## INTRODUCTION

Epilepsy is a highly prevalent neurological disorder (*1, 2*). Arising from diverse etiologies including gene mutations and environmental insults, the unifying characteristic of the epilepsies is the emergence of spontaneous, recurrent seizures (SRS). Anticonvulsants aimed at controlling seizures are the current first-line treatments for epilepsy. No current treatments have been shown to prevent or reverse disease progression; most patients remain medicated for life (*7*). Further, despite introduction of 12 new antiseizure drugs in the last 20 years, 30% of patients are drug resistant, a rate that has remained constant since 1850 (*4, 8*). Finally, with some exceptions (*9, 10*), long-term use of many anti-epileptic drugs is associated with a decline in cognition (*11, 12*), highlighting the need for transient treatments with enduring effects that address the mechanisms underlying disease mechanisms, as well as seizures.

In acquired epilepsies, epileptogenesis is the process that links the eventual propagation of spontaneous seizures to an initiating traumatic insult (*3, 13, 14*). Status epilepticus (SE), a severe bout of unremitting seizures, is the most common method for modeling epileptogenesis in acquired epilepsies. SE is associated with a number of changes in the brain including molecular, cellular, and network alterations in plasticity and inflammation (*15*). Many signaling cascades exhibit a transient period of acute activation that abates within days. How these cascades respond to the onset of spontaneous seizures during chronic disease is poorly characterized, and it is unknown whether mechanisms invoked early after insult are reengaged upon disease presentation.

Antiepileptogenic therapies aim to prevent progression to spontaneous seizures through early intervention. Although many biomarkers exist to diagnose epilepsy after the onset of seizures, a lack of predictive biomarkers for epilepsy makes an antiepileptogenic approach challenging in clinical practice (*16*). Disease modifying therapy would aim to slow or reverse the progression of chronic epilepsy after spontaneous seizures are manifest. Currently, there are no FDA approved therapeutic interventions that modify disease in an enduring manner upon brief administration (*7*).

Of the studies that have profiled gene changes in the epileptic brain, nearly all observe robust activation of inflammatory pathways that is quenched within days of SE (*17–23*). In the brain, inflammation may be triggered by glial cells (*24*) as well as neurons (*25*). Others have used anti-inflammatory drugs with varying degrees of success to suppress seizures in human patients (*26–29*), and some drugs targeting neuroinflammation in rodents have shown antiepileptogenic potential (*30–33*). However, the endogenous mechanisms that regulate neuroinflammation post-SE – acutely and chronically – are poorly understood. Recently, we applied a systems approach to a transcriptomic consortium dataset that incorporated gene expression data across multiple laboratories and models of epilepsy (*15*). Through this approach, we identified the histone methylase Enhancer of Zeste Homolog 2 (EZH2) as a potential driver of gene changes in dentate granule cells post-SE (*21*). EZH2 underwent a prolonged induction after SE across epilepsy models. Transient pharmacological inhibition of EZH2 post-SE worsened epileptic phenotypes, suggesting that EZH2 induction plays a protective role in epileptogenesis, and prompting the studies herein. In addition to EZH2, this analysis predicted Signal Transducer and Activator of Transcription 3 (STAT3) to be a driver of gene activation across models of SE.

Herein, we show that an inflammatory gene network centered around STAT3 arises across neuronal and non-neuronal cell populations acutely after SE. STAT3 itself is activated robustly and quenched in an EZH2-dependent manner within days of SE. This is followed by a resurgent activation with spontaneous seizures in the chronic period, accompanied by astrogliosis and microglial activation. Inhibiting the first wave of JAK/STAT signaling has no antiepileptogenic effect. However, briefly targeting the second wave of STAT3 activation in chronic epilepsy with the JAK inhibitor CP690550 results in an enduring, disease modifying suppression of spontaneous seizures, restoration of spatial memory, and rescue of neuropathological alterations.

## RESULTS

### Neuronal EZH2 induction protects against exacerbated epileptic phenotypes

We previously identified the histone methyltransferase EZH2 as a potential driver of gene repression post-SE in rat dentate granule cells and human TLE. EZH2 protein was robustly induced over a 10-day window post-SE in rats and mice across models of epilepsy. EZH2 induction was protective because pharmacological inhibition resulted in accelerated disease progression (*21*). To determine the role of EZH2 in epilepsy, we crossed EZH2 floxed mice (*34*) with Syapsin1 Cre drivers (*35*) to delete EZH2 in mature neurons (EZH2nHom). Neuronal specificity of Cre expression was confirmed via crossing Syn1Cre with *flox*-STOP-*flox*-tdTomato mice and immunostaining (data not shown). Staining for EZH2 confirmed reduced induction of neuronal EZH2 in EZH2nHom versus wildtype (WT) mice post-SE (Fig. S1A-D).

To test the impact of neuronal EZH2 deletion on chronic disease, we monitored behavioral spontaneous seizures in FVB WT and EZH2nHom mice by video from weeks 5-7 post-SE. EZH2nHom mice exhibited 2.5-fold greater seizure frequency compared to WT mice (unpaired t-test; p<10^−4^) (Fig. S1E). Mutant mice also had more severe Racine scale seizures than WT mice (K-S test; D:0.833, p<0.05) (Fig. S1F). These data corroborate previously published results (*21*) supporting the hypothesis that neuronal EZH2 induction post-SE is a protective response.

### EZH2 tempers activation of the JAK/STAT pathway acutely post-SE

To begin to understand the mechanism behind EZH2-mediated protection, we performed bulk RNAseq on whole hippocampal tissue from saline- and kainate-treated (4d. post-SE) WT and EZH2nHom mice. Of the 10556 expressed genes, 6241 were found to be differentially expressed at an adjusted p<10^−2^ (Likelihood Ratio Test) across the four conditions (saline, kainate, EZH2nWT, EZH2nHom). We used a community-based network approach to group genes into clusters. Leiden clustering generated 12 distinct groups (Fig. 1A). The grey, green, orange, gold, brown, skyblue, and purple clusters were generally repressed after SE, whereas the red, blue, indigo, yellow, and black clusters were generally increased. Upregulated genes were associated with inflammation, translation, post-translational modification, and glycogen metabolism, while downregulated genes were associated with cell proliferation, RNA processing, neurotransmitter release, and excitatory neurotransmission (Fig. S2A). We focused on the Black cluster because these 439 genes were induced with SE in WT mice but manifested an exaggerated induction with SE in EZH2nHom mice. We reasoned that these were genes whose expression is induced but tempered in the presence of EZH2 post-SE, and which undergo unfettered induction in the absence of EZH2.

**Fig. 1.**
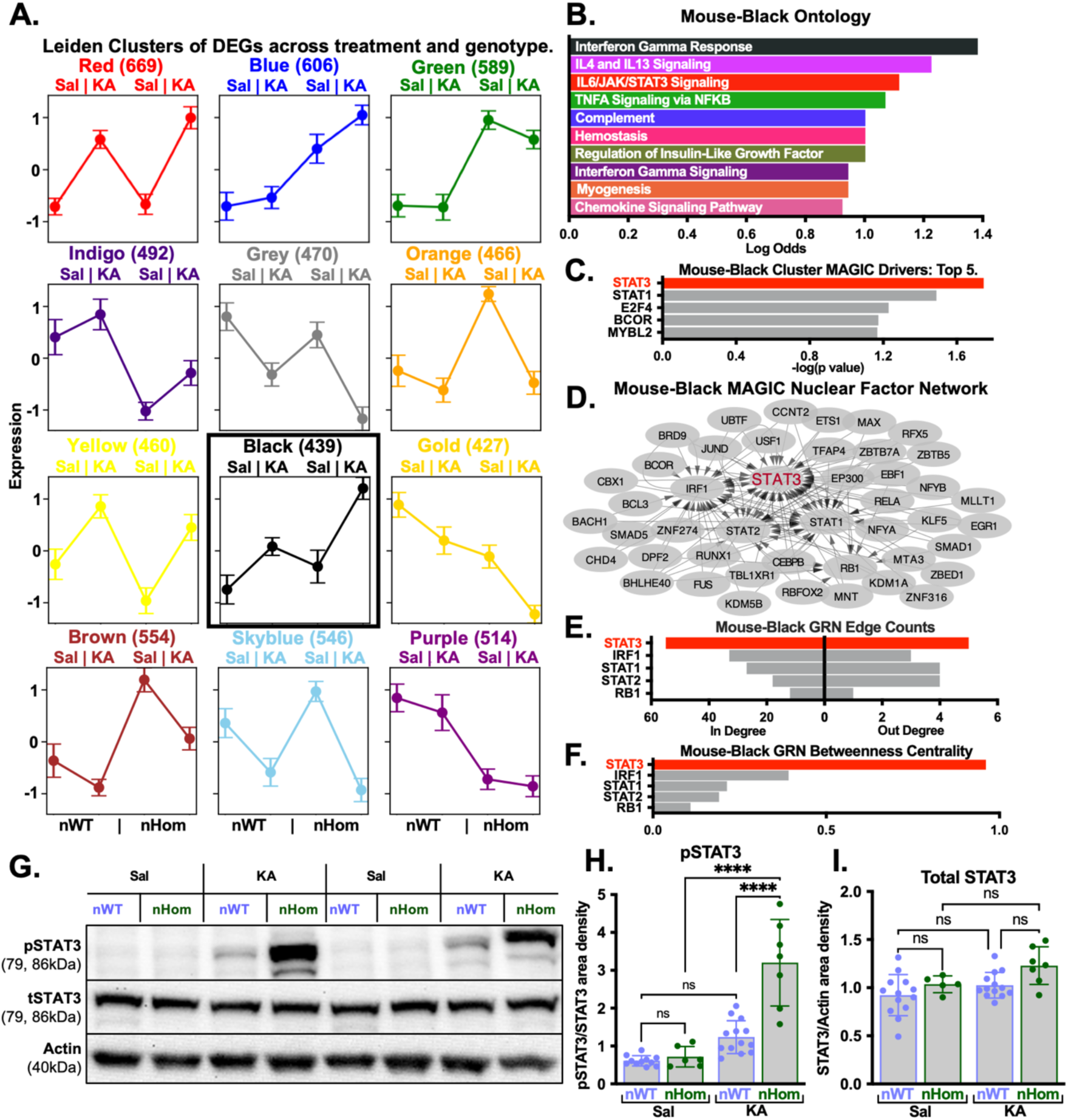
STAT3 drives hyper-activation of pro-inflammatory gene networks acutely post-SE in EZH2 knockout mice. **(A)** Black Leiden cluster of 439 genes, one of twelve generated from 6241 differentially expressed genes (fold change >1.2x, adjusted p<0.01) across saline, kainate, EZH2nWT, and EZH2nHom conditions (see Fig. S3A). **(B)** Ontological analysis of genes in the Black cluster highlighted inflammatory processes. MAGIC was used to determine which Factors control the expression of Black cluster genes. **(C)** Top 5 Factors driving expression in the Black cluster sorted by –log(adjusted p value). **(D)** Positive MAGIC hits were projected as a Gene Regulatory Network (GRN). **(E, F)** Factors were ranked by in degree and out degree counts, and betweenness centrality. **(G)** Hippocampi were harvested on day 4 post-SE from EZH2nWT and nHom mice for western blot analysis. **(H)** pSTAT3 was not altered by day 4 post-SE in EZH2nWT mice but was induced 4.5-fold (p<10^−4^) in EZH2nHom mice. **(I)** Total STAT3 levels were not altered by treatment or genotype on day 4 post-SE (2-way ANOVA with Tukey’s correction; n=5-13). *(**Seizure Model:** Systemic*

To understand the nature of genes comprising the Black cluster, we performed an ontological analysis (Fig. 1B) *(see Methods)*. The 439 Black cluster genes were compared to gene lists associated with terms in the KEGG, Reactome, Biocarta and Hallmark databases. The resulting ontology network (see Fig. S3) highlighted the innate immune response (NFkB, Interferon Gamma, Complement) and inflammation (IL6 Jak Stat3, TNF signaling) amongst others. The term ‘IL6 Jak Stat3’ was of particular interest because our previous work highlighted STAT3 as a driver of gene activation across models of epilepsy (*21*), and others have identified a pathological role for the JAK/STAT pathway in epileptogenesis (*25, 30, 36–39*). To determine which transcription factors and cofactors (hereafter together referred to as ‘Factors’) control expression of Black cluster genes, we used MAGIC (*40*) (Fig. 1C, S2B for all clusters) *(see Methods)*. STAT3 was the highest scoring Factor predicted to drive gene expression in the Black cluster. When projected as a Gene Regulatory Network (GRN), such that an arrow projects from Factor X to Factor Y if the gene encoding Factor Y is a transcriptional target of Factor X, STAT3 stood out as a central node (Fig. 1D) *(see Methods)*. Thus, STAT3 had the highest in-degree, out-degree, and total edge count (Fig. 1E) of any Factor in the network. Finally, we assessed Betweenness Centrality (*41*) to determine which Factors might be critical for information flow in the GRN. STAT3 had the greatest Betweenness Centrality of any Factor (Fig. 1F).

Acute activation of the JAK/STAT pathway post-SE in WT mice has been reported by others (*21, 37*). We observe induction of total STAT3 in neurons on day 1 post-SE (Fig. S4A). To test for EZH2-dependent changes in JAK/STAT signaling, we performed western blot analysis on hippocampal lysates from 4d. post-SE WT and EZH2nHom mice. By day 4 post-SE, phosphorylation of STAT3 at Tyr705 (pSTAT3) was diminished in WT mice compared to naïve controls, whereas EZH2nHom mice exhibited sustained pSTAT3 induction (2-way ANOVA; 4.5-fold, p<10^−4^) with no changes to total STAT3 (Fig. 1G-I). STAT3 hyperactivation was accompanied by induction of other JAK/STAT pathway components (Fig. S4B-E). These results demonstrated an EZH2-dependent suppression of STAT3 activation within days post-SE.

We considered the possibility that STAT3 hyperactivation in EZH2nHom mice was a response to worsened disease progression rather than deletion of EZH2 *per se*. Therefore, to test the hypothesis that EZH2 is required to temper STAT3 activation independent of network activity, we transitioned to an *in vitro* model. shRNA was used to knock down EZH2 in the Neuro2A (N2A) neuronal cell line (Fig. S5A). We observed robust diminution of EZH2 in knockdown cells (shEZH2) (unpaired t-test; 3.7-fold, p<10^−4^) (Fig. S5B). This was associated with a marked increase in both total STAT3 and especially pSTAT3 levels (2.4-fold p<10^−4^) (Fig. S5C-D). This result suggested that EZH2 suppresses STAT3 activation in a cell-autonomous manner without circuit level alterations.

### STAT3 drives gene expression across hippocampal cell types

To identify the factors most important for driving gene expression across cell types after seizures, we performed single nucleus RNAseq (snRNAseq) on 4 saline and 5 pilocarpine samples 4 days post-SE (Fig. 2A-B). Each sample was individually sequenced. To determine which Factors were most important for controlling gene expression across cell types after seizures, we used Deseq2 and MAGIC. Deseq2 was used to define differentially expressed gens (DEGs) for each cell type (Fig. 2C). Briefly, reads were summed for each cell type in each sample to provide a pseudo-bulked read per cell type per sample. Deseq2 was performed on this sample x gene matrix for each cell type *(see Methods)*. MAGIC was performed on the Deseq2 output for each cell type, and MAGIC outputs were analyzed using the network metrics applied in Fig.1.

**Fig. 2.**
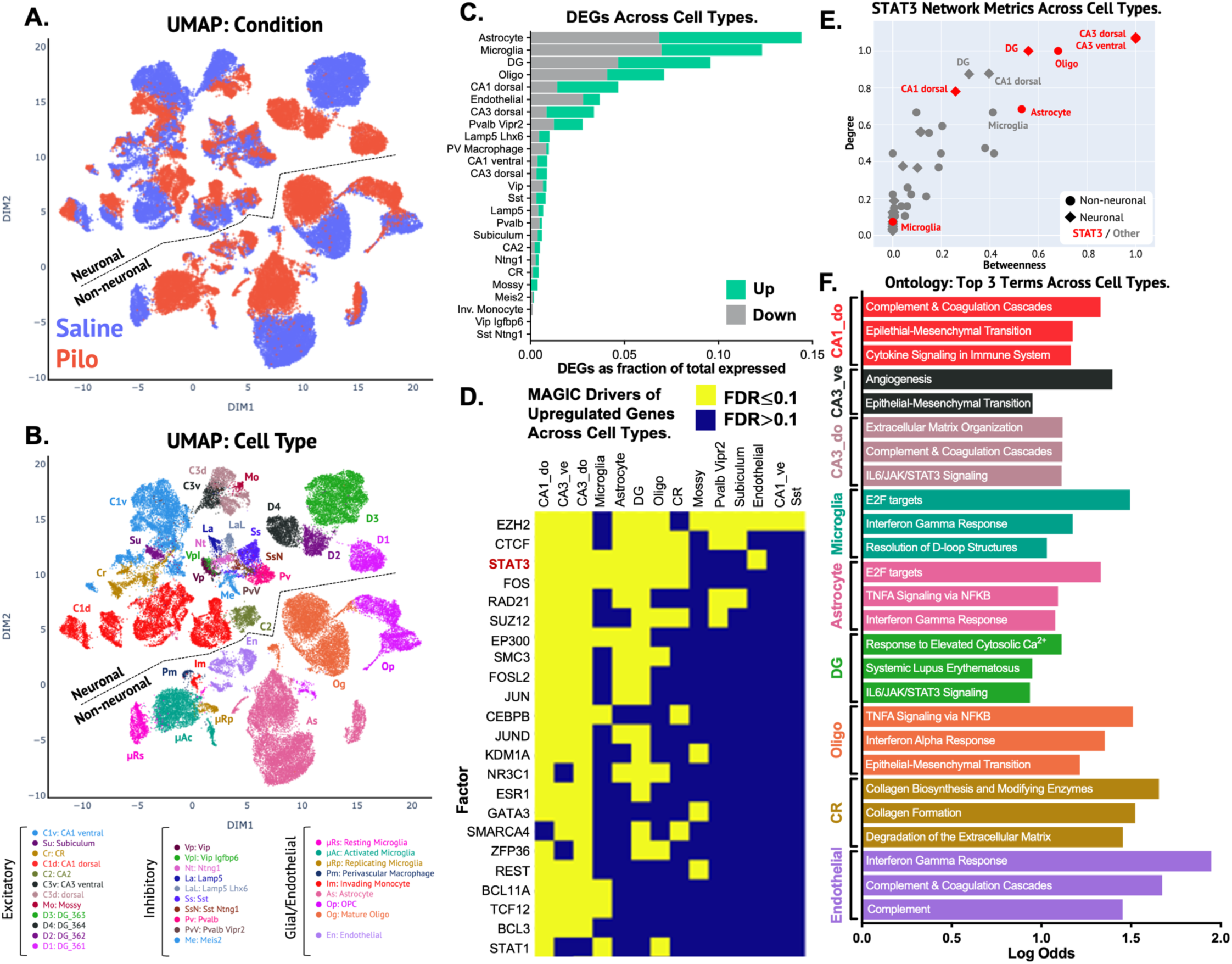
STAT3 drives gene changes across hippocampal cell types after seizures. On day 4 post-SE, 9 hippocampi harvested from saline or pilocarpine-treated mice were flash-frozen for single nuclei RNAseq (snRNAseq) (saline=5, pilocarpine=4). **(A-B)** UMAP projections display condition **(A)** and cell type **(B)**. Neurons cluster in the upper half of the UMAP, and non-neuronal cells cluster in the lower half. **(C)** The stacked bar graph shows differentially expressed genes (DEGs) across cell types, displayed as % of total expressed genes. **(D)** The heat map displays MAGIC calls for upregulated genes across cell types, with Factors ordered by the number of cell types they influence, from greatest to least. STAT3 was predicted to drive gene expression in CA1 dorsal, CA3 dorsal, and CA3 ventral pyramidal cells, microglia, astrocytes, dentate granule cells, oligodendrocytes, CR neurons, and endothelial cells. **(E)** Positive MAGIC calls were projected as GRNs for each cell type, and network metrics were tabulated for each Factor. STAT3 is the most connected (degree) and most influential Factor for controlling information flow (Betweenness) in the GRNs for CA3 dorsal and ventral pyramidal cells, oligodendrocytes, dentate granule cells, and astrocytes than any other Factor. **(F)** Ontological analysis for the cell types in which STAT3 was called as a driver was performed. Analysis highlighted tissue remodeling in neurons (CA1_do, CA3_ve, CA3_do, CR) and non-neuronal cells (Oligo), neuroinflammatory terms (IFN gamma and alpha responses, complement and coagulation, IL6/JAK/STAT3 signaling) across all cell types, DNA damage responses in microglia and astrocytes (E2F targets, Resolution of D-loop structures), and calcium signaling in DG neurons. *(**Seizure Model:** Pilocarpine)*

STAT3 was among the top three Factors controlling upregulated genes across the greatest number of cell types, including CA1 dorsal, CA3 dorsal, and CA3 ventral pyramidal cells, microglia, astrocytes, dentate granule cells, oligodendrocytes, Cajal-Retzius (CR) neurons, and endothelial cells (Fig. 2D). By the network metrics of edge count and betweenness centrality, STAT3 was the most influential Factor in the GRNs of CA3 dorsal and ventral pyramidal cells, oligodendrocytes, dentate granule cells, and astrocytes (Fig. 2E). Ontological analysis was performed on the cell types in which STAT3 was called as a driver (Fig. 2F). Among others, the analysis highlighted neuroinflammation (IFN gamma and alpha responses, complement and coagulation cascades, IL6/JAK/STAT3 signaling) across nearly every cell type. Overall, this transcriptomic analysis identified STAT3 as a central factor driving neuroinflammation across neuronal and non-neuronal cell types.

### JAK/STAT signaling is reignited in chronic epilepsy

To ascertain whether the mouse Black cluster has a counterpart in human epilepsy, we used whole transcriptome data from 129 ante-mortem human temporal lobectomy patients (*21, 42*). We used k-means clustering (*43*) to generate 10 clusters from 7658 genes co-expressed across all human samples (brown, blue, purple, skyblue, grey, orange, indigo, green, yellow, red). To test whether any of these human clusters corresponded to the mouse Black cluster, we performed Fisher exact tests to measure the degree of overlap among all mouse and human clusters (Fig. S6A-C).

The human Brown cluster showed the greatest overlap with the mouse Black cluster (Fisher exact; Odds: 4.79, p=1.19×10^−20^, 65 gene overlap) (Fig. 3A). Ontological analysis highlighted terms broadly similar to the mouse Black cluster such as inflammation (Cytokine Receptor Interaction/Complement/Innate Immune System) and interferon signaling (Interferon Gamma Response) (Fig. 3B, S6D for all best overlaps). Mirroring results of the mouse Black cluster, STAT3 had the highest in-degree, out-degree, and betweenness centrality of any Factor in the human Brown gene regulatory network (Fig. 3C-E).

**Fig. 3.**
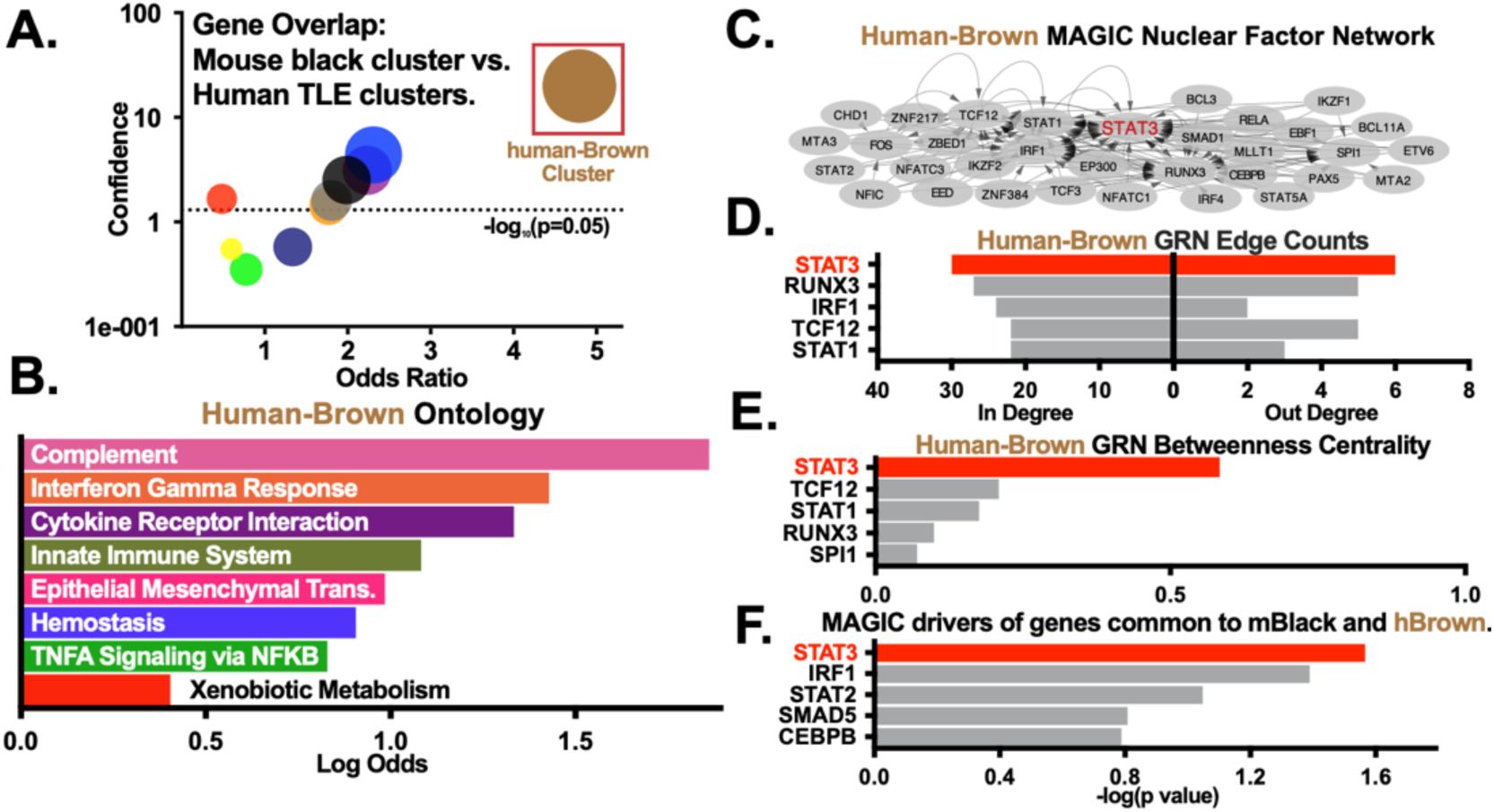
Pro-inflammatory genes activated in acute mouse epileptogenesis overlap with a module of genes in chronic human temporal lobe epilepsy. **(A)** Comparison of 10 human TLE k-means gene clusters to the mouse Black cluster by Fisher Exact test with the Benjamini Hochberg correction for multiple comparisons. Plotting -log(adjusted p) versus odds ratio against the mouse Black cluster showed that the human Brown cluster is the most similar to the mouse Black cluster (Fisher exact; Odds:4.79, p(adj)=1.2×10^−20^). Dotted line shows p=0.05. **(B)** Ontological analysis of genes in the human Brown cluster highlighted inflammatory processes. **(C)** MAGIC GRN for Factors driving the human Brown cluster. **(D-E)** Factors were ranked by in degree, out degree, and betweenness centrality. **(F)** STAT3 was the top driver predicted by MAGIC to drive expression of the 65 genes common to the mouse Black and human Brown clusters. *(**Seizure Model:** Systemic kainate)*

MAGIC called STAT3 as the highest scoring Factor driving expression of genes common to the mouse Black and human Brown clusters (Fig. 3F, S6E for all best overlaps). Given that the mouse Black cluster is from samples harvested 4 days post-SE and the human Brown cluster is from chronically epileptic humans, this data suggests that STAT3 influences disease both in acute epileptogenesis and chronic epilepsy.

Following status epilepticus, robust activation of the JAK/STAT pathway diminishes within days (*21, 37*), yet MAGIC called STAT3 as a top factor driving gene expression in the chronically epileptic human Brown cluster. Therefore, we hypothesized that STAT3 activation reignites in chronic epilepsy. To test this, we collected hippocampi from mice 1 day, 4 days, and 14 weeks after systemic kainate (Fig. 4A). On day 1 post-SE, pSTAT3 was induced 11-fold over naïve levels (2-way ANOVA; p<10^−4^), but levels were diminished by day 4 post-SE (Fig. 4A,C-D). Rapid induction and abatement of JAK/STAT signaling within days of SE was also observed in the pilocarpine model (Fig. 4B, E). After the onset of spontaneous seizures, we observed reactivation of STAT3 at 14 weeks post-SE (7.3-fold, p<10^−3^) (Fig. 4A,C-D). When pSTAT3 levels quantified across 12, 14, and 20 weeks post-SE were compared to seizure burden and frequency, we observed robust positive correlations (**Burden:** r:0.60, p<10^−3^; **Frequency:** r:0.59, p<10^−3^) (Fig. 4F-G). Other STAT isoforms were also induced in chronic epilepsy alongside STAT3, although EZH2 induction was notably absent (Fig. S7A-E).

**Fig. 4.**
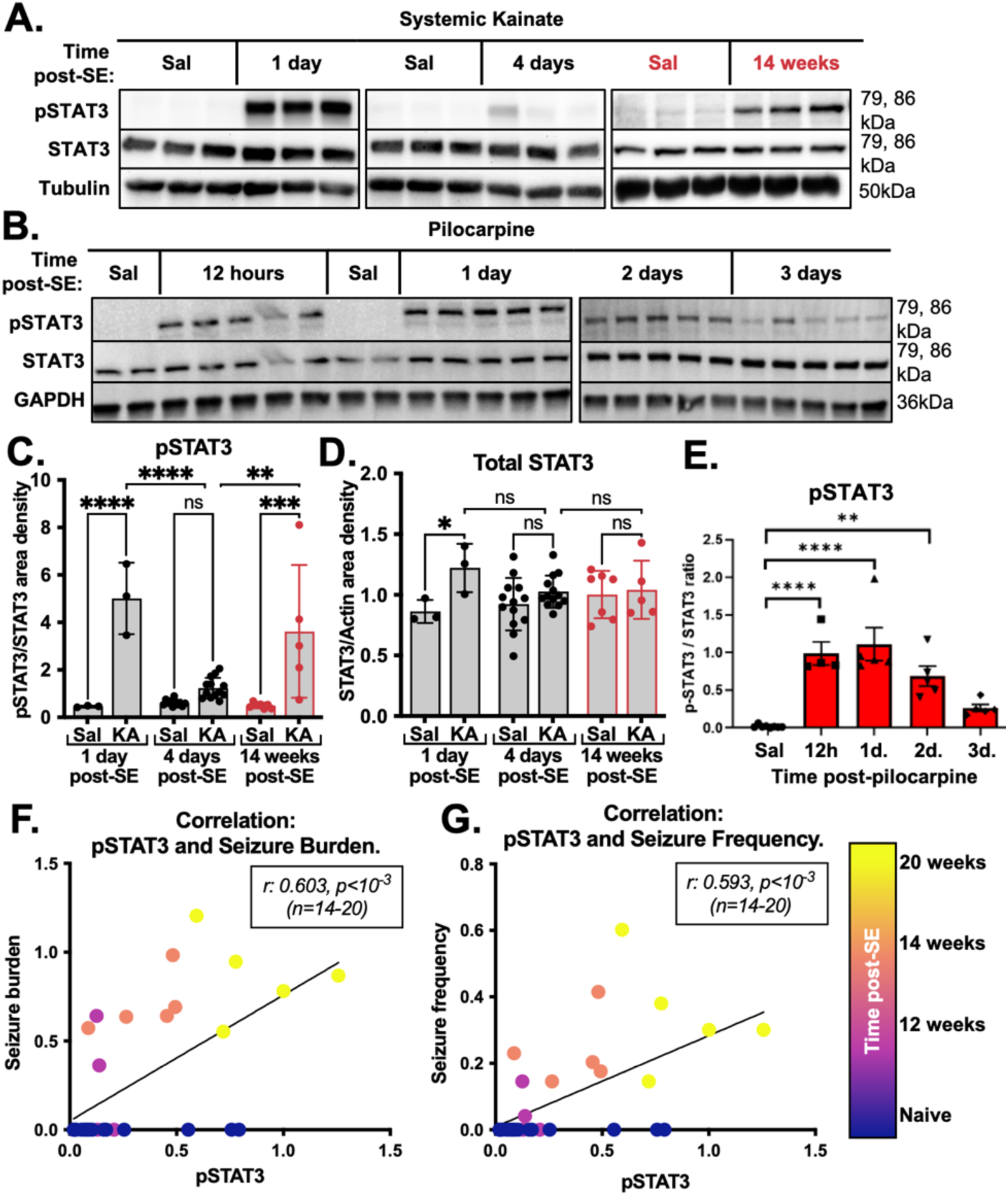
JAK/STAT signaling is transiently induced and quenched across models of SE before reactivation in chronic epilepsy. **(A)** Representative western blots for pSTAT3, total STAT3, and tubulin in saline- and kainate-injected mice at 1 day, 4 days, and 14 weeks post-SE. **(B)** Representative western blots for pSTAT3, total STAT3, and GAPDH in saline- and pilocarpine-injected mice at 12 hours, 1, 2, and 3 days post-SE. **(C-D)** In the systemic kainate model, pSTAT3 was induced 11-fold at 1 day and 7.3-fold at 14 weeks, but not induced 4 days post-SE. Total STAT3 was induced 1.4-fold on day 1 post-SE, but not 4 days and 14 weeks post-SE (2-way ANOVA with Tukey’s correction; n=3-13). **(E)** In the pilocarpine model, pSTAT3 was induced up to 48h. post-SE, but diminished by 72h. post-SE (1-way ANOVA with Tukey’s correction; n=4-8). **(F)** Seizure frequency, **(G)** burden, and pSTAT3 levels across naïve and chronically epileptic mice at 12, 14, and 20 weeks post-SE were subjected to a log(1+p) transformation. Pearson correlations (n=14-20) were performed. pSTAT3 levels were positively correlated with seizure frequency (r: 0.603, p<10^−3^) and burden (r: 0.593, p<10^−3^) in the week before harvest. (Daily seizure burden = (# of seizures * mean duration (s.))/Recording hours)) *(**Seizure Model(s):** Systemic kainate, Pilocarpine; **Seizure detection:** Video recording)*

Reactivation of JAK/STAT signaling in chronic epilepsy corroborated the overlap between the acute mouse Black and chronic human Brown clusters. We find JAK/STAT signaling is activated in neurons and glia after SE, abates within days, but undergoes reignition with spontaneous seizures.

### JAK1 is the most abundant isoform expressed across hippocampal cell types

To our knowledge, all attempts to pharmacologically modulate the JAK/STAT pathway in epilepsy have been conducted proximal to SE. Although mechanistically enlightening, this temporal window lacks a clinical application in the absence of predictive biomarkers. Further, acute intervention fails to prevent the onset of spontaneous seizures or cognitive decline as shown by others (*30, 36, 39, 44, 45*) or in our own hands (Fig. S8A-F). Instead, we asked whether we could target JAK/STAT signaling during the window of reignition in chronic epilepsy. To identify an appropriate target for pharmacological inhibition, we returned to the snRNAseq dataset described in Fig. 2. We hypothesized that the STAT3-upstream JAK kinase(s) expressed most abundantly across cell types would be the target most likely to evoke robust seizure suppression.

For every cell type, we calculated the base mean expressions of JAK1, JAK2, JAK3, and TYK2, where base mean is equal to the average number of reads across all samples regardless of condition. The values for each cell type were depicted by a dot plot (Fig. 5A, quantified in 5B-D) and UMAPs (Fig. 5E-H). JAK1 was the most abundantly expressed isoform across conditions in all cell classes (2-way ANOVA; **Excitatory:** F:13.5, p<0.01; **Inhibitory:** F:40.06, p<10^−4^; **Non-neuronal:** F:6.9, p<0.05).

**Fig. 5.**
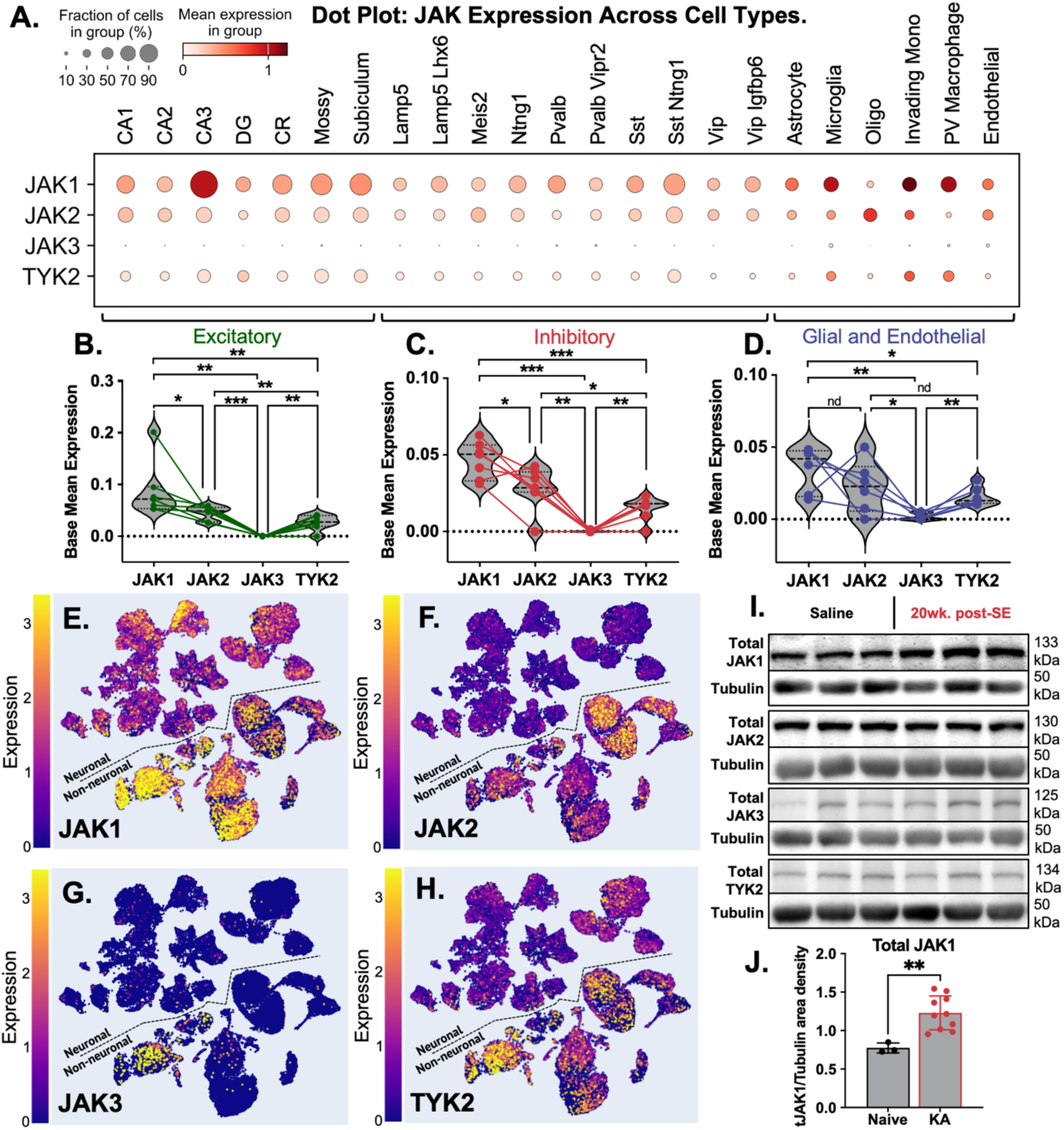
JAK1 is the most abundant isoform expressed across hippocampal cell types. **(A)** JAK isoform expression across cell types in the snRNAseq dataset described in Fig. 2 was visualized as a dot plot, where dot size describes the fraction of cells expressing the isoform and color corresponds to the mean expression in each cell class. Base Mean expression was quantified across saline and pilo samples in each cell type (e.g., CA1) for excitatory, inhibitory, and glial/endothelial cell classes and analyzed by 1-way ANOVA (BKY correction). JAK1 is the most abundantly expressed JAK isoform expressed across saline and 4d. post-SE hippocampi in **(B)** excitatory neurons (F:13.5, p<0.01), **(C)** inhibitory neurons (F:40.1, p<10^−4^), and **(D)** non-neuronal cells (F:6.9, p<0.05). **(E-H)** UMAPs were generated depicting the normalized (log(1+p)) expression of **(E)** JAK1, **(F)** JAK2, **(G)** JAK3, and **(H)** TYK2 across cell types. **(I)** Hippocampi harvested from 20wk. post-SE mice and naïve controls were subjected to western blotting. **(J)** JAK1 was induced 1.6-fold in chronically epileptic mice compared to naïve controls (unpaired t-test; p=0.0005), but JAK2, JAK3, and TYK2 were not (see Fig. S9 for quantitations). *(**Seizure Model:** Pilocarpine)*

To bridge the gap between acute epileptogenesis and the proposed drug delivery in chronic epilepsy, we assessed JAK expression via western blot, collecting hippocampi from mice 20 weeks post-kainate (Fig. 5I-J). Total JAK1 was induced 1.6-fold over naïve controls (unpaired t-test; p<10^−3^), but total JAK2, JAK3, and TYK2 were not (Fig. S9A-C). In summary, JAK1 was the most abundantly expressed across cell types days after insult, and it was induced alongside STAT3 reignition in chronic epilepsy.

### CP690550 enduringly suppresses electrographic and behavioral spontaneous seizures in chronically epileptic wildtype mice across models

To screen for potential drugs that could suppress STAT3 activation and reignition, we took advantage of the constitutive STAT3 activation in shEZH2 cells described in Fig. S5. We incubated shNT and shEZH2 N2A cells with either WP1066 (an analogue of the JAK2 inhibitor AG490 (*46–48*)) or CP690550 citrate (a small molecule inhibitor of JAK1/3 kinase activity (*5, 6, 49*)) (Fig. S10A-B). WP1066 did not alter pSTAT3 in shEZH2 cells up to 10μM, the highest concentration tested (Fig. S10C-D). In contrast, 0.5μM CP690550 citrate reduced pSTAT3 levels in shEZH2 cells to that of vehicle-treated shNT cells (2-way ANOVA; 2.3-fold inhibition, q<0.01) (Fig. S10E). These results suggest that JAK1/3 activity is required for pSTAT3 hyperactivation.

In parallel, we performed RNAi experiments in shNT and shEZH2 cells, transfecting with siRNA against JAK1, JAK2, JAK3, and a nonsense transcript (NT). This resulted in four conditions for each experiment: shNT:siNT, shNT:siJAKx, shEZH2:siNT, shEZH2:siJAKx (Fig. S10F-H). JAK knockdown was successful across all experiments (Fig. S10I-K), and JAK1 was the only isoform induced in shEZH2:siNT cells compared to shNT:siNT cells (2-way ANOVA; 1.2-fold, q<0.01) (Fig. S10I). All experiments showed induction of pSTAT3 with EZH2 knockdown, but only JAK1 RNAi succeeded in rescuing STAT3 hyperactivation back to near shNT:siNT levels (2-way ANOVA; 2-fold, q<0.05) (Fig. S10L), although shEZH2:siJAK3 cells trended towards reduced pSTAT3 (Fig. S10N). In all experiments, total STAT3 was unchanged across conditions (Fig. S10O-Q). These data support the hypothesis that JAK1 is required for STAT3 hyperactivation in N2A cells. These *in vitro* experiments concurred with our findings that JAK1 is induced in chronically epileptic WT mice and expressed ubiquitously across cell types under basal conditions and 4 days post-SE. Although we observe trends towards both JAK3 induction in chronic epilepsy (Fig. S9B) and reduction of pSTAT3 in the presence of JAK3 siRNA in N2A cells (S10N), these findings support selection of CP690550 (a JAK1/3 inhibitor) for use in animals. We therefore tested whether CP690550 treatment would reduce seizure burden when administered to chronically epileptic mice.

Daily CP690550 treatment in the chronic period alleviated seizure severity (Wilcoxon paired; W:-36.0, p<0.01) and reduced seizure frequency by 80% (1-way ANOVA; p<10^−4^) in the EZH2nHom mice described in Fig. S1 (Fig. S11A-C). We considered the possibility that CP690550 may be an anticonvulsant; we measured seizure threshold using both flurothyl and PTZ (pentylenetetrazole) (*50, 51*) in mice pre-treated with vehicle or CP690550 thirty minutes before testing. This time was chosen because peak plasma levels for CP690550 are observed 30 to 60 minutes after oral administration (*52*). CP690550 pre-treatment had no effect on seizure threshold (Fig. S11D-E).

Given that CP690550 treatment during the chronic period reduced seizure frequency in EZH2nHom mice to below wildtype levels (Fig. S11B) and the finding that EZH2 is not induced in chronic epilepsy (Fig. S7), we hypothesized that treatment during the chronic window would be effective in WT mice.

Wildtype mice were made epileptic via intrahippocampal kainate injections. An example of two spontaneous seizures is shown in Fig. 6C. In a paradigm involving one week of daily vehicle (i.p.) administration initiated 10 days after SE induction, followed by an additional week of either vehicle or CP690550 injections (15mg/kg, i.p.) under continuous EEG recording (Fig. 6A), CP690550 treatment during the chronic period profoundly suppressed spontaneous seizures. Both daily seizure frequency (mean events) and daily seizure burden ((# of events x mean duration (s))/total recording hours) were recorded. CP690550-treated mice showed a 13-fold reduction in seizure frequency (Kolmogorov-Smirnov test; D:0.91, p<10^−3^) and 8-fold reduction in burden (Kolmogorov-Smirnov test; D:0.71, p<0.01) over 1 week of recording (Fig. 6D-E). CP690550 treatment robustly reduced both spontaneous seizure frequency and burden in epileptic wildtype mice. Additionally, while all animals experienced seizures during vehicle treatment (Fig. 6B), 80% of mice were seizure-free by the third day of CP690550 treatment (Fig. 6F-G).

**Fig. 6.**
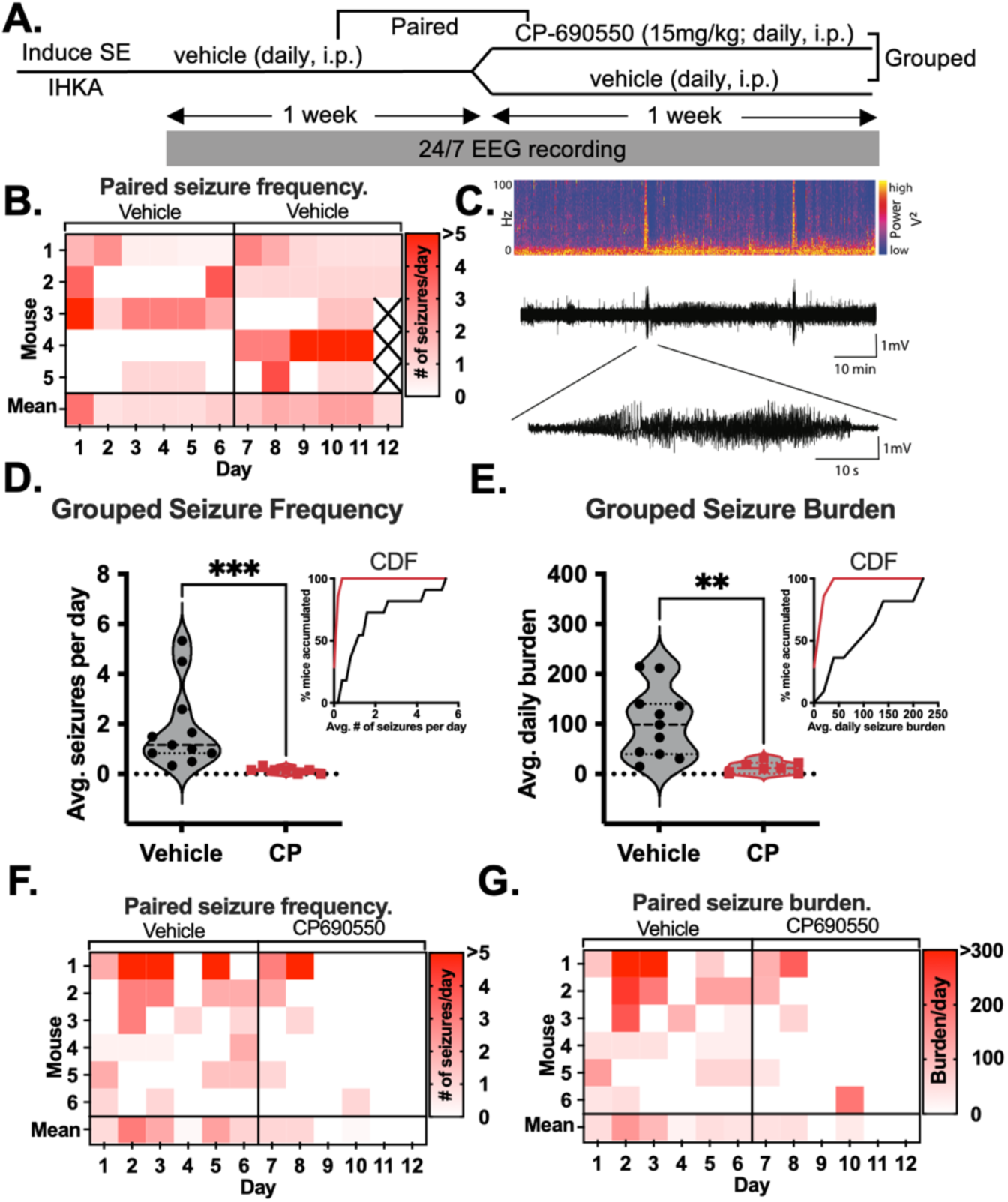
CP690550 profoundly suppresses spontaneous seizures in epileptic wildtype mice. **(A)** SE was induced via IHKA and seizures monitored by 24/7 EEG recording. Two weeks after IHKA, vehicle was administered daily for one week followed by one week of daily vehicle or CP690550 treatment. **(B)** Vehicle-treated mice had seizures across both weeks of the experiment. Seizure burden and frequency were measured for 11 instead of 12 days for mice with boxes marked by “X.” **(C)** Example power spectrum, trace, and seizure. **(D-E)** Grouped comparisons for seizure frequency and burden were displayed as violin plots and cumulative frequency distributions (bin centers are marked). CP690550 treated mice showed a 13-fold reduction in seizure frequency and an 8-fold reduction in seizure burden (Kolmogorov-Smirnov test; D:0.9091 (**Frequency**), D:0.7083 (**Burden**); n=5-12). **(F-G)** Heat maps display paired data for seizure frequency and burden. Except for one seizure observed on day 10, CP690550 eliminated seizures after two days of treatment. (Daily seizure burden = (# of seizures * mean duration (s.))/Total recording hours)) *(**Seizure Model:** IHKA; **Seizure detection:** 24/7 EEG)*

Next, we tested the therapeutic effect of CP690550 in the systemic kainate model and adopted video-based behavioral seizure monitoring. These experimental modifications aimed to generalize the efficacy of CP690550 across epilepsy models and laboratories, minimize potential confounding effects introduced by headmount implantation and associated inflammation, and prioritize clinically relevant seizure endpoints. Baseline seizure frequency and burden were recorded from weeks 8-10 post-SE (Fig. 7A).

**Fig. 7.**
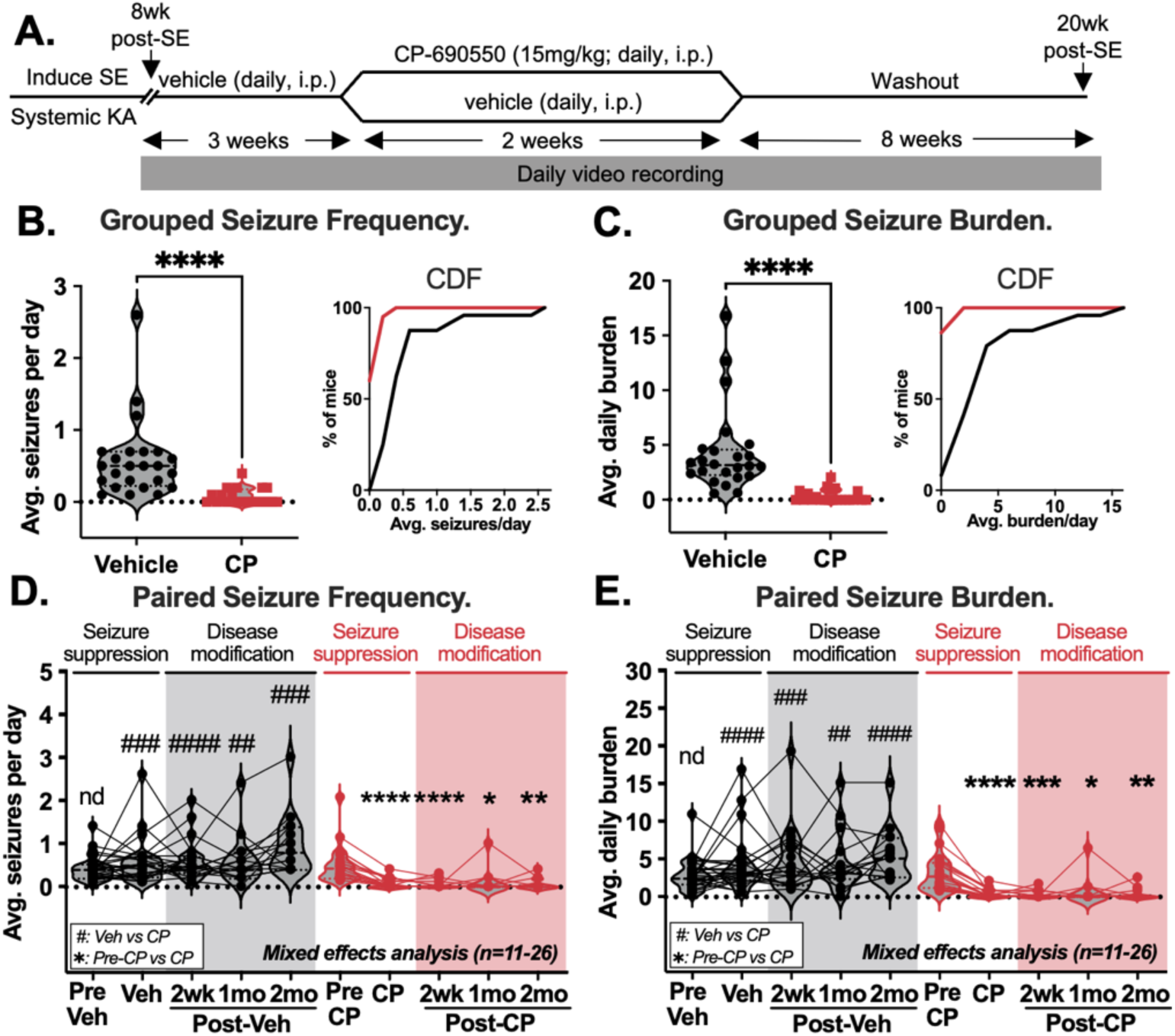
CP690550 enduringly suppresses spontaneous seizures in epileptic wildtype mice at least 2 months post-washout in the systemic kainate model. **(A)** SE was induced via systemic kainate, and behavioral seizures monitored by video from weeks 8-20 post-SE. Vehicle was administered from weeks 8-10 post-SE, followed by daily vehicle or CP690550 treatment weeks 11-12. After drug washout, video recording continued weeks 13-20 post-SE. **(B-C)** Seizure frequency and burden during treatment were displayed as violin plots and cumulative frequency distributions (bin centers are marked). Drug responders showed profound reductions in seizure frequency and burden during CP690550 treatment (Kolmogorov-Smirnov test; **Frequency:** 8-fold, D:0.7, p<10^−4^, **Burden:** 12-fold, D:0.87, p<10^−4^; n=20-24). Paired seizure frequency **(D)** and burden **(E)** before treatment, during treatment, 2 weeks, 1 month, and 2 months post-washout were displayed as violin plots with non-responders omitted (see Fig. S11, Table S1) (✱: CP-treated compared to pre-CP, #: vehicle compared to CP-treated) (Mixed effects 2-way ANOVA with BKY correction; n=11-26). 2 weeks post-CP690550, mice showed a 12-fold reduction in frequency (q<10^−4^) and a 18-fold reduction in burden (q<10^−3^) compared to 2 weeks post-vehicle; 12 out of 16 mice (75%) were seizure-free during this time. At 2 months post-washout, drug-treated animals continued to exhibit a 13-fold reduction in seizure frequency (q<0.01) and a 14-fold reduction in seizure burden (q<0.01) compared to vehicle-treated mice (n=11-15). (Daily seizure burden = (# of seizures * mean duration (s.))/Recording hours)) *(**Seizure Model:** Systemic kainate; **Seizure detection:** Video recording)*

Twenty-five mice were treated with 10 doses of CP690550 during weeks 11-12 post-SE and were recorded daily for seizures. Twenty mice (80%) responded to treatment (i.e., exhibited a seizure frequency and burden reduction of at least 50%). Drug responders exhibited an 8-fold reduction in seizure frequency and a 12-fold reduction in seizure burden (Fig. 7B-C) (Kolmogorov-Smirnov test; **Frequency:** D:0.7, p<10^−4^, **Burden:** D:0.87, p<10^−4^). Even incorporating the non-responders (i.e., the 5 animals that displayed <50% reduction in seizure frequency and burden), we observed a 3-fold reduction in frequency and a 4-fold reduction in burden (Fig. S12 and Table S1 for analysis with all animals).

We aimed to assess the durability of CP690550’s effect on seizures. To do this, we withdrew drug from responders and continued monitoring seizure frequency and burden from 13-20 weeks post-SE. CP690550 eliminated seizures completely in 65% of drug responders during treatment. Two weeks after drug withdrawal, 75% of mice were seizure-free (2-way ANOVA; **Frequency:** 12-fold, q<10^−4^, **Burden:** 18-fold, q<10^−3^) (Fig. 7D-E). Some mice were harvested on the final day of treatment (12 weeks post-SE) and at two weeks post-washout (14 weeks post-SE) for experiments described in Figures 4 and 9 *(see Methods)*.

Seizure suppression persisted to the end of recording. One month post-CP690550, one drug responder showed a relapse; 8 out of 10 mice were seizure-free. Two months after the last dose of CP690550, we observed a 13-fold reduction in seizure frequency and a 14-fold reduction in seizure burden compared to vehicle-treated mice (2-way ANOVA; **Frequency:** q<0.01; **Burden:** q<0.01). Together, these data suggest that CP690550 treatment may be a potent, novel therapeutic strategy for lasting seizure suppression.

### CP690550 reverses cognitive comorbidities

We asked whether chronically epileptic mice treated with CP690550 showed improvement of epilepsy-associated deficits in spatial memory. We tested working and short-term spatial memory at three time points: 1-2 weeks prior to SE, during CP690550 treatment (12 weeks post-SE), and 2 weeks after drug washout (14 weeks post-SE) (Fig. 8A).

**Fig. 8.**
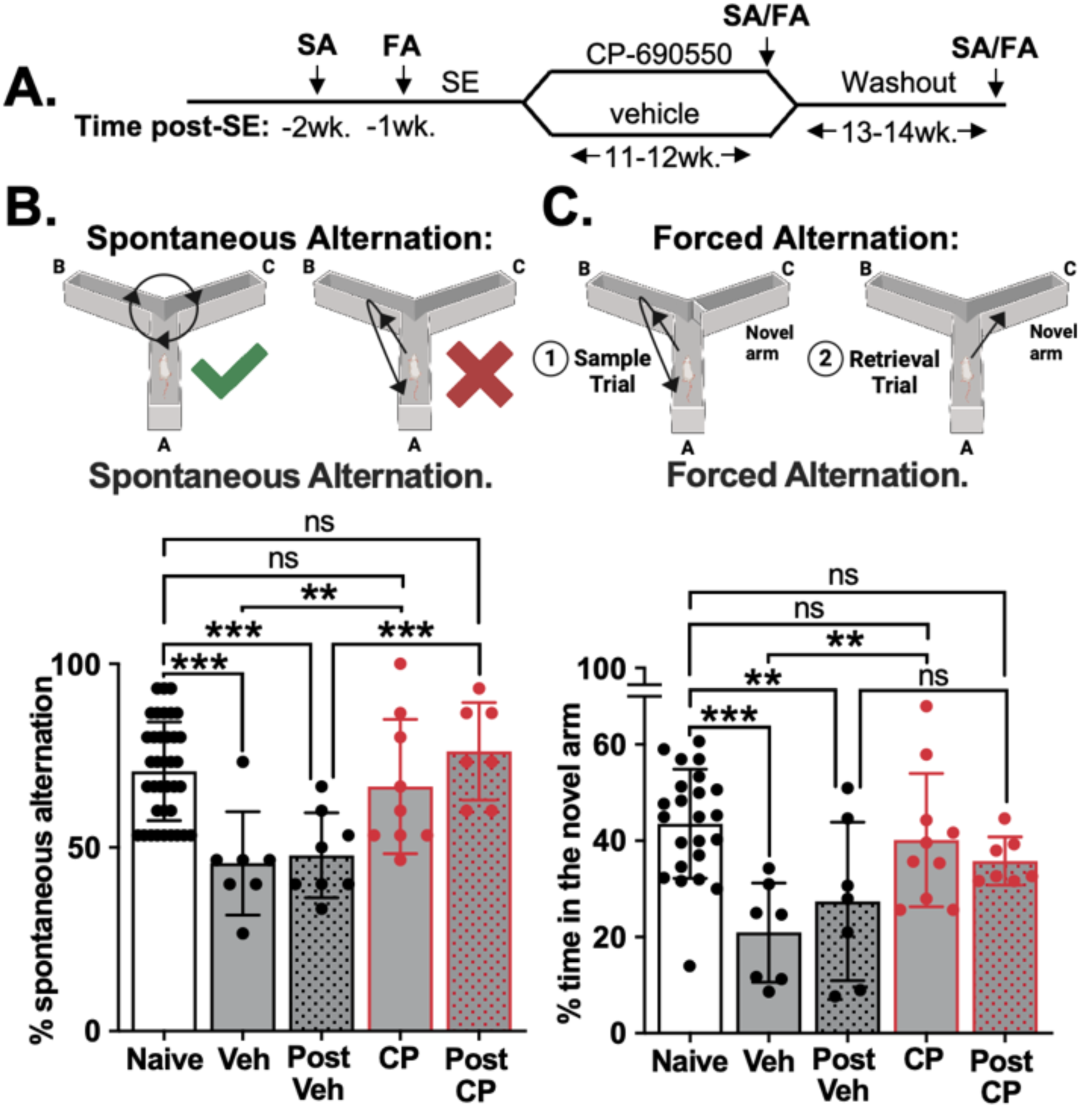
CP690550 rescues working and short-term memory deficits in chronically epileptic mice. **(A)** Outline of experimental approaches for subsequent panels. **(B)** Spatial working memory was severely impaired in untreated chronically epileptic mice; SA was reduced 35% from baseline at week 12 post-SE and 32% at week 14 post-SE. Mice treated with CP690550 showed a 31% rescue in SA at week 12 post-SE and a 37% rescue at week 14, compared to vehicle-treated mice (1-way ANOVA with BKY correction; n=7-37). **(C)** Spatial short-term memory was likewise impaired in untreated mice; FA was reduced 52% from baseline at week 12 post-SE and 37% at week 14. In contrast, mice treated with CP690550 showed a 48% rescue in FA at week 12 post-SE compared to vehicle-treated mice (1-way ANOVA with BKY correction; n=7-23). Daily treatment with CP690550 rescued both SA and FA to levels indistinguishable from naïve mice both during treatment and after drug washout. *(**Seizure Model:** Systemic kainate)*

First, we asked whether CP690550 could rescue working spatial memory. To do this we measured spontaneous alternation (SA) in a previously described Y-maze testing paradigm (*32, 53, 54*). Mice were placed in the starting arm of the maze facing the end of the arm and allowed to explore freely for 8 minutes. Sequential arm choices during exploration of a three-arm Y-maze are considered an indicator of spatial working memory because the innate curiosity of a healthy mouse promotes exploration of the most novel arm (i.e., the arm least recently visited) (*53, 54*). Distance traveled during each test was measured to ensure deficits in SA post-SE were not due to mobility impediments (Fig. S13A). Chronically epileptic mice exhibited a 35% reduction in SA compared to naïve animals (1-way ANOVA; q<10^−3^) (Fig. 8B). In contrast, daily CP690550 treatment rescued spatial memory by 31% (q<0.01), restoring performance to naïve levels (Fig. 8B).

Short-term memory requires that information collected during an experience be consolidated and accessible shortly thereafter (*55*). Thus, we asked whether CP690550 treatment could rescue short-term memory, as measured by forced alternation (FA) tests in a Y-maze as previously described (*7, 53*). As with working memory, chronically epileptic mice showed profound deficits in short-term spatial memory, with time spent in the novel arm reduced by 52% at 12 weeks post-SE (1-way ANOVA; q<10^−3^) compared to naïve mice (Fig. 8C). Mice treated daily with CP690550 exhibited a 48% increase (q<0.01) in time spent in the novel arm (Fig. 8C).

To assess the durability of CP690550’s impact on cognition, we withdrew drug after week 12 post-SE. Two weeks after withdrawal, SA and FA tests were again performed. It should be noted that in naïve mice, repeated testing does not alter results for spontaneous or forced alternation, nor does CP690550 treatment in naïve mice (Fig. S13B-E). At 2 weeks post-washout, CP690550-mediated restorations in working memory persisted with a 37% increase in percent spontaneous alternations compared to vehicle-treated mice (1-way ANOVA; q<10^−3^) (Fig. 8B). Short-term memory as measured by time spent in the novel arm remained indistinguishable from naïve animals two weeks after drug withdrawal, consistent with an ongoing protective effect of CP690550, although no significant difference was noted between post-vehicle and post-CP690550 mice, likely due to large variance in the former group (Fig. 8C). Together, these data show that targeting reignition of JAK/STAT signaling in chronic epilepsy enduringly suppresses both seizures and cognitive comorbidities.

### CP690550 alleviates epilepsy-related histopathological alterations

Profound seizure suppression observed with CP690550 treatment prompted us to assess the impact of this drug on histopathological markers of epilepsy. Hippocampi were collected from naïve mice (Fig. 9A), mice on the final day of treatment with either vehicle (Fig. 9B) or CP690550 (Fig. 9C), and mice 2 months post-vehicle (Fig. 9D) or -CP690550 (Fig. 9E) (n=1-2). Sections were probed for DAPI, total STAT3 (*I-V*), GFAP (*VI-X*), IBA1 (*XI-XV*), and P2Y12 (*XVI-XX*).

**Fig. 9.**
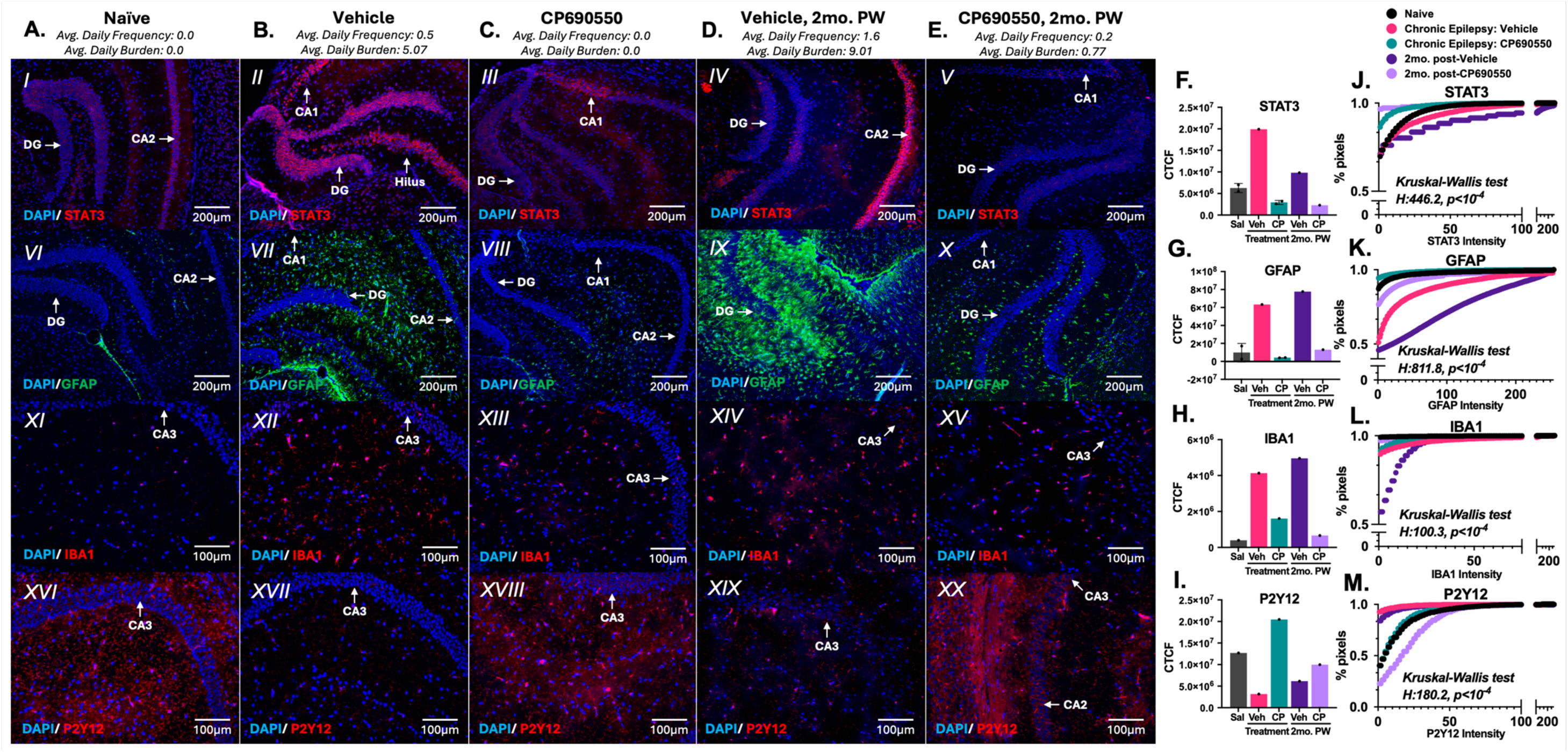
CP690550 treatment enduringly suppresses astrogliosis and microglial activation at least 2 months post-washout. SE was induced via systemic kainate, and seizures monitored by video from 8-20 weeks post-SE. Hippocampi were harvested from **(A)** naïve mice, **(B-C)** mice 12 weeks post-SE (final day of CP690550 or vehicle treatment), or **(D-E)** 20 weeks post-SE (2 months post-vehicle or -CP690550) were subjected to immunostaining for total STAT3 (I-V), GFAP (VI-X), IBA1 (XI-XV), P2Y12 (XVI-XX), and DAPI (n=1-2). Average daily seizure frequency and burden prior to harvest is listed for each animal shown in panels A-E. Representative were quantified by corrected total cell fluorescence **(F-I)** and by intensity **(J-M)** (see Methods). Intensity histograms were converted to cumulative curves, and distributions across treatment groups were compared via Kruskal-Wallis tests (Dunn’s correction). **(F, J)** At the end of drug treatment, total STAT3 expression was induced in vehicle-treated chronically epileptic mice (II) compared to naïve (I) and reduced in CP-treated mice (III). 2 months after drug withdrawal, total STAT3 expression was persistently induced in chronically epileptic mice (IV) compared to naïve (I) and reduced in mice 2mo. post-CP (V). **(G, K)** GFAP **(G, K)** and IBA1 **(H, L)** were both reduced in CP-treated (VIII, XIII) mice at the end of drug treatment compared to vehicle-treated mice (VII, XII) and remained suppressed 2mo. after withdrawal of CP690550 (X, XV). **(I, M)** P2Y12, a marker of microglial homeostasis, was downregulated in chronically epileptic mice during vehicle treatment (XVII) and 2mo. post-vehicle (XIX). CP690550 treatment rescued P2Y12 expression during treatment (XVIII); this effect persisted at 2mo. post-CP690550 (XX). *(**Seizure Model:** Systemic kainate; **Seizure detection:** Video recording)*

Representative images were quantified both for corrected total cell fluorescence (Fig. 9F-I) and fluorescence intensity (Fig. 9J-M) *(See Methods)*. Total STAT3 induction was used as a proxy for activated STAT3, for lack of an antibody against pSTAT3 that can be used for immunostaining in mouse brain. Chronically epileptic mice showed a profound induction of total STAT3 in neurons compared to naïve animals (p<10^−4^), while CP690550 treatment reduced total STAT3 expression (p<10^−4^) to levels indistinguishable from naïve mice (Kruskal-Wallis test with Dunn’s correction) (Fig. 9F,J; *I-III*). Two months after drug was withdrawn, STAT3 expression remained suppressed in post-CP690550 mice compared to post-vehicle (p<10^−4^) (Fig. 9F,J; *IV-V*).

CP690550-mediated reductions in neuronal STAT3 expression were further accompanied by changes in astrogliosis and microglial activation. Both GFAP and IBA1 were induced in chronically epileptic mice (**GFAP:** p<10^−4^; **IBA1:** p<10^−4^) and reduced by CP690550 treatment (**GFAP:** p<10^−4^; **IBA1:** p<10^−4^) (Fig. 9G-H,K-L; *VI-VII,XI-XIII*). Two months post-CP690550, astrogliosis was still reduced compared to post-vehicle mice (p<10^−4^), but microglial activation was not (p=0.19) (Fig. 9G-H,K-L; *IX-X,XIV-XV*).

P2Y12 is a purinergic receptor selectively expressed on resident microglia (*56*) with reported roles in microglial homeostasis and neuroprotection (*57–59*), and its downregulation has been observed in association with both neurodegenerative disease (*60*) and seizures (*61*). Herein, P2Y12 showed downregulation in chronically epileptic vehicle-treated mice compared to naïve (p<10^−3^), but levels were rescued with CP690550 treatment (p<10^−4^) (Fig. 9I,M; *XVI-XVIII*). Two months after the last dose of drug, P2Y12 levels remained downregulated in post-vehicle mice compared to post-CP690550 (p<10^−4^) (Fig. 9I,M; *IX-XX*). Chronically epileptic mice treated with CP690550 exhibited rescue of histopathological changes and lasting suppression of STAT3 expression months after drug withdrawal.

## DISCUSSION

We show that rapid induction of JAK/STAT3 signaling after SE is quenched by the chromatin modifier EZH2, but reignites in chronic epilepsy. Targeting this resurgence with CP690550 reverses cognitive comorbidities and extinguishes astrocytic and microglial activation in an enduring manner. We provide evidence that CP690550 is a novel, disease-modifying therapeutic agent for durable seizure suppression, cognitive restoration, and neuropathological rescue across epilepsy models. CP690550 (Tofacitinib) is an FDA-approved orally available JAK1/3 inhibitor indicated for moderate-to-severe rheumatoid arthritis and other inflammatory diseases. Work presented here supports the use of Tofacitinib to reduce seizure burden in epilepsies with a substantial neuroinflammatory component. Because its mechanism of action is different from those of conventional antiseizure drugs, its antiseizure effects are predicted to be additive or synergistic with those of other epilepsy drugs.

### EZH2 induction is protective after neurological insults

After status epilepticus, EZH2 protein is induced in hippocampal pyramidal and dentate granule neurons across models of epilepsy (*21*). Previously, we reported that treatment of mice with the SAM-competitive EZH2 inhibitor UNC-1999 exacerbated spontaneous seizures (*21*), and here we find that deletion of EZH2 in mature neurons has the same effect. Induction of Polycomb group (PcG) proteins has been studied in association with other brain insults. For example, sublethal ischemic insults lead to induction of Polycomb in mouse cortical neurons (*62*). Further, sevoflurane (a volatile anesthetic that exerts positive allosteric modulation on GABA_A_ receptors (*63*) and antagonizes NMDA receptors (*64*)) causes protective induction of EZH2 and H3K27Me3 to attenuate hypoxic-ischemic brain injury and improve post-injury behavioral assessments in rats. These improvements are reversed when sevoflurane is given alongside the EZH2 inhibitor GSK126 (*65*). These studies, along with our findings, point to a potentially general protective role for EZH2 after CNS traumas.

Weaver syndrome, a congenital disorder characterized by partial PRC2 loss-of-function due to EZH2 missense mutations, is typified by bone overgrowth, cognitive deficits, and seizures, among other disease features (*66–68*). For experiments utilizing EZH2nHom mice, we observed no weight differences, reduced seizure thresholds, or altered spatial memory in 6 week-old naïve EZH2nHom animals compared to EZH2-sufficient littermate controls (EZH2nWT) (Fig. S1G-I). Work from others has revealed that even frank EZH2 deletion in chondrocytes cannot recapitulate Weaver syndrome phenotypes (*69*). Furthermore, Weaver syndrome involves germline EZH2 mutations affecting all cell types, unlike our neuron-specific deletion, which suggests our observations in EZH2nHom mice are distinct from Weaver pathology and likely represent a trauma-specific protective role of neuronal EZH2 (*70*).

### EZH2 tempers activation of JAK/STAT signaling after status epilepticus

EZH2 deletion in neurons results in large transcriptomic changes after SE (Fig. 1). The mouse Black gene cluster comprises genes induced after SE in EZH2nWT mice and hyper-induced in EZH2nHom mice after SE. This pattern suggests that EZH2 tempers expression of genes in this cluster because its deletion results in their unfettered activation. Ontological analysis of the mouse Black cluster produces terms associated with innate immunity, neuroinflammation, and the JAK/STAT pathway. Network analysis places STAT3 in the center of the mouse Black GRN. Increased activation of STAT3 at Tyr705 as a result of EZH2 inactivation has also been observed in a transformed murine cell model of early T cell precursor acute lymphoblastic leukemia; EZH2 inactivation leads to enrichment of documented STAT3 target genes and exaggerated pSTAT3 induction with IL6 stimulation (*71*). STAT-upstream receptors and kinases are also regulated by EZH2. In prostate cancer EZH2 represses IFNGR1 to dampen JAK2/STAT1 signaling, and its depletion restores expression of IFN target genes and pSTAT1 (*72*). The exact mechanism by which EZH2 tempers JAK/STAT signaling during epileptogenesis is a current focus of the lab.

### STAT3 drives gene expression across cell types after seizures

Bulk RNAseq is inherently limited by the inability to resolve cell type-specific gene alterations. As a result, meaningful changes in underrepresented cell types (e.g., classes of inhibitory neurons, invading monocytes, etc.) may be occluded by highly represented populations (e.g., astrocytes, CA1 pyramidal neurons, etc.). Thus, we took advantage of the resolution and unprecedented statistical power of an snRNAseq dataset comprising hippocampi from four naïve mice and five mice harvested four days after pilocarpine SE. Samples were sequenced separately. In so doing, we aimed to answer the question of whether STAT3 was a driver of gene changes across cell types, or in just one or two overrepresented populations. Our rationale was that any putative target for disease modification ought to be a principal driver of gene expression across neuronal and non-neuronal cell types, such that interventions might interrupt the cross-talk between populations that contributes to disease propagation (*73–76*). Indeed, we found that STAT3 drives gene changes across neuronal and glial cells after pilocarpine-induced SE, and this Factor is the most connected and most crucial hub for gene regulatory networks in astrocytes, dentate granule cells, oligodendrocytes, microglia, and CA3 pyramidal cells after seizures. These cell populations exhibit some distinct functions (e.g., DNA damage responses in microglia), but they are unified by terms related to neuroinflammation and innate immunity, underlining and corroborating the predicted, programmatic role STAT3 occupies in each population.

### JAK/STAT signaling is reactivated in chronic epilepsy

Analysis of human TLE samples identifies a cluster of genes similar to the STAT3-driven mouse Black cluster; the human Brown cluster is a group of genes associated with innate immunity and neuroinflammation (Fig. 3). Like the mouse Black cluster, the most influential driver of gene expression is STAT3. The robust overlap between clusters of genes, one from frankly epileptic human samples and another from rodents in an early epileptogenic period, suggests a couple of insights. Firstly, STAT3-mediated epileptogenic mechanisms identified in mice are likely clinically relevant to the human population. Secondly, STAT3 drives gene changes both acutely after insult and in chronic epilepsy; this is supported by the identification of STAT3 as a driver of gene expression, its transient activation acutely post-SE, and its reactivation in chronic rodent epilepsy (Fig. 4). Furthermore, the failure to observe EZH2 induction in chronic epilepsy (Fig. S7) suggests its protection is limited to acute epileptogenesis, as animals with intact neuronal EZH2 still go on to develop spontaneous seizures, and the same genes hyper-induced with EZH2 deletion after SE are widely co-expressed across drug refractory human TLE patients.

STAT3 phosphorylation is robustly induced on day 1 post-SE and diminished by day 4 post-SE (Fig. 4). Its subsequent resurgence in chronic epilepsy occurs at least as early as 12 weeks post-SE and persists to at least 20 weeks post-SE (Fig. 4,9). It remains unclear whether reignition of JAK/STAT signaling precedes or follows the onset of spontaneous seizures. This would be difficult to assess because latency to spontaneous seizure onset after chemoconvulsant-induced SE varies widely by model, mouse strain, and delivery method (*77, 78*). In the kainate model of epilepsy (*79, 80*), others have observed latencies to spontaneous seizures ranging from 5-30 days for the IHKA model and 10-30 days for the systemic kainate model (*77*). Detection method also matters because electrographic seizures typically arise earlier than “clinical” motor seizures. Mean latency to electrographic seizures ranges from 5-13 days post-SE (*81–83*), while mean latency to behavioral seizures is more variable, ranging from 10-77 days post-SE (*77, 81–84*).

If STAT3 reactivation precedes the onset of spontaneous seizures, this suggests that inflammatory JAK/STAT signaling itself may drive the onset of seizure activity, and putative mechanisms by which this may occur have been explored by the Brooks-Kayal and Russek groups (*25, 85*). If STAT3 reactivation occurs after the onset of spontaneous seizures, then the enduring effect on seizures that we observe with CP690550 treatment suggests that ongoing inflammation in chronic epilepsy is required for seizures. In either case, STAT3 reactivation is targetable and potently disease modifying.

### JAK/STAT signaling is a potent therapeutic target in chronic epilepsy

Pharmacological inhibition of STAT3 prior to or during SE blunts, but does not prevent onset of spontaneous seizures (*30*). Though mechanistically enlightening, this approach lacks clinical utility in the absence of reliable disease biomarkers. Attempts to genetically ablate the STAT3 SH2 domain (*36*) plateaued disease progression but did not reduce seizure burden, and the impact on comorbidities was unclear due to differences between naïve wildtypes and naïve STAT3-SH2 knockout mice (*36*). Harnessing our snRNAseq dataset, we were able to identify JAK1 as the most abundantly expressed JAK isoform across all cell types (Fig. 5), and show that JAK3 is induced in microglia after pilocarpine SE. Further *in vitro* studies led to the selection of the JAK1/3 inhibitor, CP690550, for pharmacological intervention *in vivo* (Fig. S10). With delivery during the chronic period and subsequent rescue of the exacerbated seizure phenotype in EZH2nHom mice (Fig. S11), we identified the target of a novel endogenous protective mechanism invoked after seizures. In this mechanism, EZH2 acts to temper JAK/STAT signaling days after insult, and EZH2 loss leads to worsened disease progression; the molecular underpinnings of this mechanism are a current focus of the laboratory.

Moving to WT mice, we find inhibiting the first wave of JAK/STAT signaling with CP690550 has no effect on spontaneous seizures and epilepsy-associated memory decline (Fig. S8). Targeting the second wave of STAT3 activation with the JAK inhibitor CP690550 results in an enduring suppression of spontaneous seizures, restoration of spatial memory, and reductions in multiple histopathological markers of inflammation (Fig. 6-9). Whether seizure suppression and cognitive restoration are *caused by* reduced inflammation is an open question. Administration of chemoconvulsant agents like kainic acid or pilocarpine recapitulate neuropathological and electrographic features of human temporal lobe epilepsy (*86*). However, we have not assessed the effectiveness of this agent in models which replicate the features of genetic, developmental, TBI-or tumor-related epilepsies. It remains to be seen whether CP690550 would be effective for epilepsies in which inflammation does not play a primary role in disease progression.

CP690550 profoundly suppresses seizures in chronically epileptic wildtype mice across models of epilepsy as measured by two different methods of seizure detection. In the systemic kainate model, CP690550 improved seizure frequency and burden by at least 50% in 80% of mice (20 out of 25 mice). Fig. S12 and Table S1 show that the potently disease modifying effect of CP690550 treatment persists even with non-responders included in the analysis. Following cessation of CP690550 treatment we observe robust disease modification for at least two months. The endurance of this protection is the focus of ongoing studies. Our data are inconclusive as to whether JAK1 or JAK3, or both, is the relevant target of CP690550.

Our finding that CP690550 can be administered transiently in the chronic period to give lasting seizure protection would largely mitigate the risk of systemic immunological effects associated with long-term use of JAK inhibitors, as in rheumatoid arthritis (RA) (*87*). RA patients may require up to 2 years of CP690550 treatment before improvements begin to manifest (*88, 89*). Although two weeks of daily CP690550 treatment in mice would be roughly equivalent to 18 months in humans by simple life-span scaling (*90*), the minimum required dosing period for extended protection has yet to be determined. For now, we find the ratio of protection time to treatment time is at least 4:1. Deaths during or after vehicle or CP690550 treatment were noted and analyzed for any treatment effects. There was no significant difference in mortality between treatment groups (Log-rank Mantel-Cox test; Chi square:0.893, p>0.05) (Fig. S12). Given the rapid suppression of seizures within 3 days of treatment (Fig. 6), future studies will seek to shorten the course of treatment, test intermittent dosing, and address the impacts, if any, of the current treatment regimen on cardiac health (*91*) and immune function (*92*).

### CP690550 restores working and short-term memory in chronically epileptic mice

We show that in addition to seizure suppression, CP690550 treatment during the chronic period alleviates epilepsy-associated deficits in spatial memory (Fig. 8). Notably, CP690550 administered acutely post-SE has no effect on short-term memory decline (Fig. S8), highlighting the importance of this second, targetable window. Cognitive decline is not only a challenge with current epilepsy therapies, but also a leading comorbidity of epilepsy itself (*93, 94*). Despite recent advances, there is a continued, critical need for disease modifying agents that address comorbid behavioral and cognitive decline(*10, 95*). The mechanism(s) linking epilepsy to comorbidities is still unclear, but inflammation is a commonality across epilepsy and co-occurring conditions such as cognitive dysfunction (*96, 97*), depression (*98*), anxiety (*99*), and schizophrenia (*100*). Indeed, others have shown that suppression of key neuroinflammatory mediators such as COX2-associated pathways (*32*) and Complement 3-dependent synaptic remodeling (*101*) rescues aspects of cognitive function in preclinical rodent models of acquired epilepsy. To our knowledge, CP690550 is the first pharmacological agent that may be administered transiently after the onset of spontaneous seizures to rescue spatial memory for weeks after drug withdrawal.

### CP690550 rescues histopathological alterations in chronic epilepsy

We assessed histopathological alterations in chronic epilepsy by performing immunofluorescence on hippocampal tissue harvested from naive, vehicle-treated, and CP690550-treated chronically epileptic mice on the final day of treatment (Fig. 9). Representative images showed a significant increase in the expression of STAT3, GFAP, and IBA1 in vehicle-treated mice, indicating heightened inflammation and gliosis characteristic of chronic epilepsy (*102–105*). STAT3 expression was most elevated in neurons but did show expression across cell types. In contrast, CP690550-treated mice exhibited a marked reduction in the levels of STAT3, GFAP, and IBA1. Interestingly, P2Y12, which was downregulated in vehicle-treated epileptic mice, was upregulated in both naive and CP690550-treated mice, suggesting a restoration of microglial homeostasis (*57–59*). Two months after drug washout, we repeated immunostaining for the same targets. Expression patterns observed at the end of treatment persisted, with CP690550-treated mice continuing to show reduced levels of STAT3 and GFAP, alongside sustained upregulation of P2Y12. These findings suggest an enduring anti-inflammatory effect with brief CP690550 treatment.

While activation of the JAK/STAT pathway in non-neuronal cells post-SE is well-characterized (*24, 38, 106, 107*), the role of STAT3 in neurogenic inflammation post-SE is less clear (*36*). Cytokines released from glial cells following seizures increase neuronal excitability (*108*), and type I interferons increase excitability in neocortical pyramidal neurons via PKC (*109*). Others report that seizure-induced activation of the JAK/STAT pathway alongside upregulation of BDNF reduces the number of gamma-aminobutyric acid (GABA) type A alpha1 receptors expressed in neurons following seizures, influencing GABA(A)R-dependent inhibition (*25*). This relationship is likely reciprocal, with the reduction of neuronal STAT3 expression influencing inflammatory markers in non-neuronal cells. For example, BDNF upregulated in neurons after seizures binds to TrkB receptors expressed by glial cells to promote reactive phenotypes (*110, 111*). By dampening STAT3 activity in neurons, CP690550 may indirectly reduce the reactivity of astrocytes and microglia, as indicated by the decreased levels of GFAP and IBA1 (Fig. 9).

### Comparison of CP690550 with other disease modifying agents

The persistence of CP690550’s impact on seizures, cognition, and histopathological alterations underscores its potential as a disease-modifying agent. Existing anti-inflammatory drugs that have been used to suppress seizures in human patients (*26–29*) or have shown antiepileptogenic potential in rodents (*30–33*) pose several drawbacks: some or all are proposed only as supplemental therapies (*26, 27*), risk worsened disease outcomes (*27*), or show only transient effects (*28*). The disease modifying properties of agents like dimethyl fumarate (DMF) (*112*) and sodium selenate (*113*) have been reported for models of acquired epilepsy. The use of senolytics has also been successful in reducing seizure frequency in a mouse model of focal cortical dysplasia type II, a cortical malformation associated with drug-resistant pediatric epilepsy (*114*).

Like CP690550, DMF is FDA-approved (*115*), and its administration in chronic epilepsy suppresses seizures during treatment and after drug washout. Whereas DMF was able to enduringly suppress seizures when administered to chronically epileptic rats, its impact on cognition with chronic administration was not tested, and administration at the time of insult, as previously highlighted, has little clinical utility. CP690550 is distinguished compared to other drugs, in that it appears to offer enduring benefits associated with reduced inflammation long after drug discontinuation. This long-term modulation of inflammatory markers is notable, as it points to a sustained impact on an underlying neuropathological process in epilepsy.

### Systemic inflammation in rheumatoid arthritis is associated with higher risk of epilepsy

CP690550 citrate is an FDA-approved drug for the treatment of rheumatoid arthritis and ulcerative colitis, and evidence points to an association between peripheral inflammatory disease and risk for developing epilepsy. Adjusted hazard ratios for developing epilepsy were compared between a cohort of 32,000 rheumatoid arthritis (RA) patients and a cohort of sex- and age-matched controls; RA patients were at a significantly elevated risk (*116*). Similarly, a bout of sepsis caused a 5.2-fold increased incidence of seizures over an eight-year follow-up period (*117*). Intriguingly, when RA patients were stratified by duration of nonsteroidal inflammatory drug (NSAID) use, those who had used NSAIDs for less than 1.5 years were at the greatest risk, while those using NSAIDs for more than 5.8 years exhibited adjusted hazard ratios for epilepsy that were lower than the non-RA control cohort.

### Conclusions

We have employed an open discovery, omics-based approach involving multiple animal models and laboratories to identify drivers of disease, paired with a concerted effort to confirm relevance to human disease. With this approach we identify EZH2 as a protective negative regulator of the JAK/STAT pathway in epileptogenesis. Our current data demonstrates that CP690550 mitigates spontaneous seizures in chronic epilepsy and is disease modifying. The cross-model, systems-based approach by which we identified JAK/STAT signaling as a target of EZH2 and selected CP690550 suggests this drug may be useful for other neurological disorders. In sum, characterization of an endogenous, EZH2-mediated, protective response across rodent models of epilepsy has highlighted a biphasic activation of JAK/STAT3 signaling: an early induction and abatement after insult, followed by reactivation with chronic, spontaneous seizures. Transiently targeting STAT3 reignition in chronic epilepsy with the FDA-approved drug CP690550 is potently seizure suppressing, memory restoring, and disease modifying.

## METHODS

### Bioinformatics

#### Brief Lay Summary of Bespoke Bioinformatic Scripts and Methods

All bioinformatic tools described herein are Python scripts that will be made available on Github.

##### Leiden Clustering Algorithm

The Leiden algorithm groups genes based on how similarly they are expressed across different conditions using a tool in the Python Scanpy package (*118*). Higher “resolution” settings create more clusters, and lower settings create fewer. Each cluster includes genes that show similar behavior, such as rising or falling expression. This clustering helps in identifying groups of genes that may be functionally related in experiments with more than 2 conditions.

##### Ontomancer

Ontomancer identifies biological functions and pathways for gene sets. It compares each gene set to gene sets associated with biological terms across ontological databases (KEGG, Reactome, Biocarta, Hallmark) from https://www.gsea-msigdb.org/gsea/downloads.jsp. For each biological term, Ontomancer tests whether there is a significant overlap (by Fisher Exact test) with the gene set and genes associated with the biological term. Results are organized in a file with each worksheet showing enriched terms from a respective database.

##### Ontomancer Network Maker

This tool creates a network of related biological terms from Ontomancer’s results. Biological terms from across databases that share genes are connected, and the strength of each connection depends on how many genes overlap. The network is then divided into “communities,” and the community takes on the name of the highest scoring term in that community. This allows ontological results from across databases to be visualized in a single image and makes it easier to see which biological processes are linked. Visualization tools are used to display these networks clearly.

##### MAGIC

MAGIC identifies transcription factors, co-factors, or chromatin modifiers that might regulate gene changes in each cluster (*40*). MAGIC uses ChIPseq data from ENCODE to see if these factors bind more strongly to genes that change in an experiment versus all the genes expressed in an experiment. Significant factors are listed with test results in a summary file, and additional files show how well each factor binds to specific gene sets.

##### MAGIC Network Maker

This tool creates a gene regulatory network (GRN) using the results from MAGIC. Factors that are predicted to drive gene changes in an experiment are projected as a network such that arrows connect Factor X to Factor Y if Factor Y is a target of Factor X. The network is viewed and analyzed in Cytoscape (*119*).

#### Bulk RNA Sequencing

##### Tissue Harvest, RNA Extraction, and Sequencing

Hippocampi were harvested from saline- and kainate-treated EZH2nWT and EZH2nHom mice on day 4 post-SE (n=4-5 per treatment/genotype). RNA was extracted from hippocampal tissue with Trizol (#1559602, Life Technologies, Carlsbad, CA), and concentration was determined by spectrometry. This was followed by a treatment with DNase (#AM2238, Invitrogen). RNA clean-up was carried with the RNeasy Mini Kit (#74104, Qiagen, Hilden, Germany). Purified samples were submitted to the Gene Expression core at the University of Wisconsin-Madison Biotechnology Center for library preparation and sequencing (https://biotech.wisc.edu).

##### Sequencing, Expression Quantification, and Differential Gene Expression

Reads were mapped to the mm39 build mouse genome and subjected to Likelihood Ratio Tests using Deseq2 (*120*). Transcripts per million were calculated and genes were classified as expressed if they had at least 3 TPM in all samples of any one of the 4 conditions (saline:WT, kainate:WT, saline:EZH2nHom, kainate:EZH2nHom). Deseq2 output was then filtered for only expressed genes and adjusted p-values were recalculated for the reduced number of genes. Genes with an adjusted p<.01 and 1.2-fold change were subjected to Leiden clustering with resolution=0.5 using the scanpy Leiden tool (*118*). The Leiden algorithm clusters genes based on how similarly they are expressed across different conditions. Genes within a cluster show similar behavior across the 4 conditions. This clustering helps to identify groups of genes that may be functionally related in experiments with more than 2 conditions.

##### Ontology, MAGIC, and Gene Regulatory Network Analysis

Ontology was performed on all clusters using Ontomancer (manuscript in preparation). Briefly, Biocarta, Hallmark, KEGG, and Reactome .gmt files were downloaded from https://www.gsea-msigdb.org/gsea/downloads.jsp. Gene sets associated with every term in the databases were collected. For each Leiden cluster, Scores (S_t_) for each term, t, were calculated as:

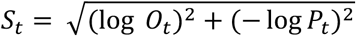

Where O_t_ and P_t,_ are the odds ratio and p values from Fisher exact tests for the overlap of Leiden cluster genes and genes for each term t. Only terms with odds ratios>1 and p values<0.05 were kept for further analysis. For each term, a gene list is generated containing the overlap of genes in the cluster and term, G_t_. A term x term similarity matrix was generated where similarity between term i and j is the Fisher exact test log odds ratio O_i,j_ for the intersection of genes in the terms G_i_ and G_j_. This was converted to a network where nodes are terms. Edge weights were calculated as:

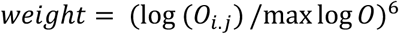

Where max log odds is the maximum log odds value across all comparisons. The 6 power was used to separate nodes for visualization. Network communities were generated using the scanpy Leiden tool with resolution=2. Each community was named after the highest scoring term in that community. Networks were visualized via Python’s networkx and Plotly tools.

MAGIC was performed on all clusters essentially as described(*40*) but with r=1, and GTFs were ignored. GTFs were defined as any ENCODE Factors containing the strings “TBP”, “POL”, “GTF”, or “TAF”.

Gene Regulatory Networks were projected using Cytoscape (*119*) using MAGIC outputs. Factors were connected to their targets with edges defined as ChIP signal extracted from the Target Folder generated by MAGIC. Target genes of Factors that were not themselves Factors were removed to generate the GRN. Network statistics were calculated in Cytoscape (*119*). ***Code for generation of Figures 2-3 will be made available on GitHub*.**

#### Single Nuclei RNA Sequencing

##### Tissue Harvest and Sequencing

To select which epileptic mice would be sent for sequencing, we determined a composite score of seizure severity during pilocarpine SE, weight loss 24 hours after SE, ability to build a nest and neurobehavioral recovery scores on day 4 after SE (See Fig. S14). The mean number of standard deviations from the mean of each of these 4 measures was calculated across all 6 saline-treated and 14 pilocarpine-treated mice, and mice were selected based on wide separation between saline and pilo groups. Both hippocampi and a small portion of neocortex overlaying one hippocampus were snap frozen and shipped to Novagene.

##### Sequencing, Preprocessing, Expression Quantification and Quality Control

Each sample was individually subjected to library preparation and Illumina sequencing. Reads were processed and mapped with Cell Ranger software (10x Genomics) and the Seurat R package at Novagene. Counts matrices for each sample were combined into an adata object in the Roopra lab and subjected to quality control, cell and gene filtering using the Scanpy package (*118*). Genes were kept that were expressed in at least 100 cells across all samples. Cells with genes numbering between 1000 and 6500 and with fewer than 5% mitochondrial genes were kept. This resulted in 64,647 cells and 21,423 genes.

For PCA analysis of samples, reads were summed across all cells for each sample to produce a pseudo-bulked sample (n=10) x gene (21,423) array. 4000 most variant genes were selected to subject samples to PCA analysis. All 5 pilo samples clustered together when projecting either the 1^st^ 2 or the 2^nd^ 2 PCA components. Four saline samples clustered together away from the pilo samples. One saline sample did not cluster with either group and was discarded (Fig. S14). This resulted in 5 pilo samples, 4 saline samples, and 58,365 cells.

##### Normalization and Clustering

Data was normalized using the scanpy preprocessing normalization tool. Highly expressed genes were omitted, and the target sum was set to 10,000 (transcripts per 10k or TPTK). Counts were converted to log(1+tptk) and highly variable genes identified with min_mean=0.0125, max_mean=3, min_disp=0.5. Genes were scaled to unit variance and values>10 standard deviations were clipped. PCA with 50 components was performed and used to generate a neighborhood graph with n_neighbors=15. Leiden clustering with resolution=0.25 yielded 27 clusters.

##### Annotations

###### Class annotation

Clusters were allocated to classes using the following markers: Inhibitory Neurons: GAD1, Excitatory Neurons: SLC17A7, Astrocytes: AQP4, Oligodendrocytes: SOX10, Endothelial cells: RGS5, Myeloid: PTPRC. Each Class were broken up into Cell Classes as follows:

###### Excitatory Neurons

The single cell dataset from Yao et al. (*121*) was converted to an anndata object and trimmed to glutamatergic neurons (CA1-ProS, CA2-IG-FC, CA3, CR, DG). DEGs/markers were identified for each subtype using the rank_genes_groups scanpy function with method=’wilcoxon’. The same was done for our Excitatory adata object using groupby=’leiden’. The top 100 Yao markers per cell type and our leiden clusters were used to perform pairwise fisher exact tests to look for overlaps between Yao cell types and our leiden clusters. The Yao cell type with highest overlap with Leiden pairs as judged by highest score (log(odds)*-log(p)) was assigned to the cluster. This yielded the following subclasses: CA1_ProS, CA1_ve, CA1_do, CA3, CR, DG. Using Hipposeq (*122*) the CA3 subclass (markers: [SPOCK1, MGAT4C, ELAVL2]) was broken up into CA3_ve, CA3_do and mossy cells using the following markers: CA3_ve: [NECAB1,COCH,ADGRA1,CPNE7], CA3_do: [CHGB,RIMBP2], mossy: [CALCRL,CALB2]. Yao et al CR cell markers were RELN, CACNA2D2, STMN1. DG markers were PROX1, PCP4, ADARB2, STXBP6.

###### Inhibitory Neurons

As with excitatory neurons, the Yao dataset was trimmed to GABAergic neurons. Rank genes groups was run on the inhibitory adata object and the Yao adata object. Fisher exact tests with the top 100 genes per group was performed for each Yao subtype and Inhibitory leiden cluster. This approach defined 10 inhibitory subtypes in our dataset: Lamp5 Lhx6, Lamp5, Meis2, Ntng1, Pvalb Vipr2, Pvalb, Sst Ntng1, Sst, Vip, Vip Igfbp6.

###### Astrocytes

AQP4 was used to define astrocytes. GFAP was used to define activated and resting states.

###### Myeloid cells

The myeloid cell class was first defined by PTPRC. The myeloid adata object was then reclustered with n_neighbors=15 and leiden resolution=0.25. This gave 12 clusters. GFAP defined activated microglia. SIGLCECH positive and PTPRC negative marked resting microglia. Cells with high TOP2A, PCNA and MKI67 defined proliferating microglia. CCR2 and CD44 positive denoted invading monocytes. MRC1 and VCAM marked Perivascular Macrophages.

###### Oligodendrocytes

The oligodendrocyte cluster was reclustered with n_neighbors=15 and leiden resolution=0.25 to give 9 clusters. Mature oligodendrocytes were marked by MOG. Progenitors (OPC in Fig. 2) were marked by VCAN.

##### Differential Gene Expression, MAGIC, and Ontology

Deseq2 was used for differential gene analysis between saline and epileptic samples for each cell class (Astrocyte, CA1, CA2, CA3, CR, DG, Endothelial, Invading_Monocyte, Lamp5, Lamp5 Lhx6, Meis2, Microglia, Mossy, Ntng1, Oligo, Perivascular_Macrophage, Pvalb, Pvalb Vipr2, Sst, Sst Ntng1, Subiculum, Vip, Vip Igfbp6). For each cell class, reads for each gene were summed within a sample such that each sample contributed a single integer value (total reads) for each gene. Thus, a sample x gene matrix was generated for every cell class. Each cell class sample x gene matrix was passed to Deseq2 and Wald tests were performed to find DEGs between saline and pilo samples. Deseq2 output was then filtered for expressed genes in each cell class. Expressed genes were defined as those genes that had at least 1 read in every sample of either condition. After filtering Deseq2 output, padj was recalculated using the lower gene number. DEGs were genes with an absolute log fold change of ≥1.2 and a padj<0.05. Deseq2 outputs for each cell class were subjected to MAGIC and ontology as for bulkseq. For single cell gene network analysis, MAGIC-derived Factor target genes for each cell class were used to generate GRNs in the Python Networkx package(*123*). Betweenness and degrees were extracted from networkx output dictionaries.

#### Human TLE expression data analysis

##### Expression Quantification, Normalization, and Clustering

GSE63808 (*42*) from GEO contained 129 transcriptomes from resected hippocampal TLE tissue. Probes with an Illumina p-value<0.05 in at least 20% of samples were retained in the analysis; the probes were consolidated to gene symbols by taking the median probe expression value per symbol. These 9076 genes formed the background list for ontological and MAGIC analyses. Expression values were normalized by the median expression value in that sample. Genes kept for further analysis exhibited variance greater than the median coefficient of variance across all samples. The resulting 7658 unique genes were subjected to k-means clustering using the k-means tools from Sci-kit learn (*124*) to divide genes into 10 clusters (brown, blue, purple, skyblue, grey, orange, indigo, green, yellow, red).

##### Overlap Across Mouse and Human Clusters

Overlap between each of the 10 human TLE clusters with the 12 mouse clusters was measured by Fisher exact test with the Benjamini-Hochberg correction for multiple comparisons. Overlap was measured by: number of genes in common, confidence (-log_10_adjusted p), and score (odds ratio x -log_10_adjusted p). For each mouse cluster, the human TLE cluster considered to have the best overlap was the one which showed the highest score. Table S3 shows the statistics from every test, as well as the gene symbols for each overlap.

##### Ontology, MAGIC, and Gene Regulatory Network Analysis

Ontological and MAGIC analyses were performed on the overlapping genes for each highest-scoring mouse/human cluster combination. For all genes in the human Brown cluster – the human TLE cluster that overlapped most robustly with the mouse Black cluster – ontological analysis was performed, and the gene regulatory network was constructed and analyzed.

### Cell culture

#### Growth conditions

Cells were grown in 5% CO_2_ at 37°C. Neuro2a cells were grown in MEM (#10-009-CV Corning, Manassas, VA) with 1.0g/L glucose, 1.5g/L L-Glutamine, and 10% fetal bovine serum (#26140-079 Gibco, Grand Island, NY).

#### Lentiviral knockdown

Stable EZH2 knockdown in Neuro2a cells for western blot was achieved using SMARTvector lentiviral delivery of shRNA per the manufacturer instructions. Puromycin selection began 48 hours after infection and maintained during cell expansion and experimentation. SMARTvector lentiviral mouse EZH2 mEF1a-TurboRFP shRNA (#V3SM7592-232015353, Dharmacon, Lafayette, CO) targeted the sequence *ATCGTAGTAAGTACCAATG*, and non-targeting mEF1a-TurboRFP control particles (#S10-005000-01) were used as an infection control. These generated EZH2 knockdown (shEZH2) and control (shNT) cells.

#### siRNA knockdown

Transient siRNA knockdowns were performed on shEZH2 and shNT Neuro2a cells. Cells were transfected with ONTARGET-plus SMARTPool siRNA (Dharmacon, Lafayette, CO) using the Lipofectamine RNAiMAX Transfection Reagent (#13778030, ThermoFisher, Waltham, MA). Cells were harvested 48 hours after transfection. Knockdowns were performed against mouse JAK1 (#L-040117-00-0005), JAK2 (#L-040118-00-0005), JAK3 (#L-040119-00-0005), and a non-targeting pool (#D-001810-10-05).

#### WP1066 and CP690550 citrate

WP1066 (#573097, Millipore, Bedford, MA) or CP690550 citrate (#4556, Tocris Bioscience, Bristol, United Kingdom) were solubilized in DMSO. shEZH2 and shNT Neuro2a cells were treated with either WP1066, CP690550 citrate, or DMSO and were allowed to incubate for one hour before protein was harvested. Cells were treated with WP1066 to a final concentration of 0μM (DMSO), 3μM, or 10μM. Cells were treated with CP690550 citrate to a final concentration of 0μM (DMSO), 0.5μM, or 5μM.

### Molecular Experiments

#### Tissue isolation and homogenization

After KA seizure induction and/or treatment with CP690550 citrate, animals were sacrificed by decapitation. Whole hippocampal hemispheres were harvested and flash-frozen in liquid nitrogen or on dry ice. Hippocampal tissue was lysed in Radioimmunoprecipitation Assay Buffer (RIPA: 50mM Tris, 150mM NaCl, 1% nonidet P-40, 0.5% sodium deoxycholate, 0.1% SDS) with mammalian protease inhibitor (1:1000, Sigma or 1860932 ThermoScientific) and phosphatase inhibitor (1:1000, Sigma or 78428 ThermoScientific). Protein from cell lines was harvested in Triton lysis buffer (3% 5M NaCl, 10% glycerol, 0.3% Triton X-100, 5% 1M Tris pH8.0) after PBS washes. Tissue was homogenized in lysis buffer by probe sonication (Fisher Scientific, Sonic Dismembrator, Model 100, Hampton, NH) on power 3 for two rounds, with 10 pulses per round. Supernatants were isolated by centrifugation and quantified using the DC Protein Assay (Bio-Rad, Hercules, CA) or BCA assay (ThermoScientific 23227). Protein extracts were stored at −80°C. 5x loading buffer (0.5mM Tris, 10% SDS, 50% glycerol, 10mM EDTA, and 1% beta-mercaptoethanol) was added to each sample to reach a 1x final concentration. Extracts in loading buffer were boiled at 95°C for 5 minutes and stored for up to one month at - 20°C until run on an acrylamide gel.

#### Western blotting

##### Systemic Kainate

Following systemic kainate, protein extracts in loading buffer were loaded at 20µg per lane and resolved by electrophoresis in hand-poured acrylamide gels with a 5% acrylamide stacking layer (125mM Tris pH6.8, 5% acrylamide, 0.01% ammonium persulfate, 0.01% SDS, 0.01% TEMED) and an 8% acrylamide separating layer (375mM Tris pH8.8, 8% acrylamide, 0.015% ammonium persulfate, 0.015% SDS, 0.08% TEMED). Gels were transferred to polyvinyl difluoride membranes (PVDF; Millipore, Bedford, MA) using Tris-glycine transfer buffer (20mM Tris, 1.5M glycine, 20% methanol). Membranes were blocked with 5% bovine serum albumin (Fisher Scientific, Fair Lawn, NJ) diluted in low-salt Tris-buffered saline (w/w TBST; 20mM Tris pH7.5, 150mM NaCl, 0.1% Tween 20) with mammalian protease inhibitor (1:2000) and phosphatase inhibitor (1:2000), or Odyssey blocking solution (LI-COR, 927-60001) for 1 hour at room temperature. Primary antibodies were diluted in the same blocking buffer and incubated with membranes overnight at 4°C. Antibodies include: EZH2 (1:1000, #5426 Cell Signaling, Danvers, MA), phospho-STAT3 (Tyr705) (1:2000, #9145 Cell Signaling), total STAT3 (1:1000, #12640 Cell Signaling), JAK2 (1:1000, #3230 Cell Signaling), STAT1 (1:1000, #14994 Cell Signaling), pS727-STAT1 (1:1000, #8826 Cell Signaling), STAT2 (1:1000, #72604 Cell Signaling), pY1007/08-JAK2 (1:500, ab32101, Abcam, Cambridge, UK), Tubulin (1:1000, #T9026 Sigma-Aldrich), and Actin (1:10,000, #691001 MP Biomedicals).

The next day, the membranes were washed three times in 1X TBST and incubated with horseradish peroxidase-conjugated goat-anti-mouse or -rabbit secondary antibodies for one hour at room temperature (1:10,000, Invitrogen #31430, #31460, Rockford, IL). Membranes were subsequently washed three times in TBST, and membranes were developed in SuperSignal West Femto ECL reagent (ThermoFisher, Waltham, MA). Bands were imaged using a ChemiDoc-It Imaging System (UVP Vision-Works, Upland, CA) and quantified using UVP Vision Works software or ImageJ. Band intensities were graphed and analyzed using Prism 9 software (GraphPad Software, La Jolla, CA).

Blots probed for pSTAT3 were stripped and re-probed for total STAT3 and tubulin as shown in Figures 1G, 4A, S4B, S5A, S7A, S8E, and S10A, B, F, G, and H. Stripping was performed by washing the blot for 10 minutes in TBST after imaging, followed by a 15 minute incubation in Restore Western Blot Stripping Buffer (Thermo Scientific, #21063) at 37°C. These conditions were validated by washing the stripped blot in TBST for 10 minutes and reimaging to ensure the secondary antibody was stripped. To ensure the primary antibody was stripped, the blot was re-blocked, incubated in secondary antibody for one hour, washed, and reimaged.

##### Systemic Pilocarpine

Following systemic pilocarpine, after perfusion with ice-cold 1xPBS, brains were carefully removed from the skull and hippocampi were dissected, frozen on dry ice and stored at −80 °C until further use. Each hippocampus was homogenized by 15 strokes in 200 µL RIPA buffer (Thermo Scientific, 89901) containing a protease (Thermo Scientific, 1860932) and a phosphatase (Thermo Scientific, 78428) inhibitor cocktail, followed by centrifugation at 10,000×g for 10 min at 4 °C. Supernatant was collected, and protein concentration was determined by BCA assay (Thermo Scientific, 23227). Twenty micrograms of protein for each sample were separated on SDS polyacrylamide gels running at 150V for 1 h, transferred at 100V to a PVDF membrane for 1 h in the cold room, and then blocked with Odyssey blocking solution (LI-COR, 927-60001) for 1 hour at room temperature. Membranes were then incubated with primary antibodies against phosphorylated-STAT3 (p-STAT3, 1:2,000, Cell Signaling, 9145) and GAPDH (1:10,000; Millipore, CB1001) overnight at 4 °C. After washing 3x with TBS-0.1% Tween-20 (TBST, VWR, 0777), membranes were incubated with the secondary anti-rabbit (IRDye800, 1:10,000, LI-COR, 925-32211) and anti-mouse (IRDye680, 1:10,000, LI-COR, 925-68020) antibodies for 1 hour at room temperature. For total STAT3 detection, membranes were then stripped with Restore blot stripping buffer (Thermo Scientific, 21059) for 15 min at room temperature. After stripping, membranes were washed 1x with TBST, blocked with Odyssey blocking solution (LI-COR, 927-60001) for 1 h at room temperature and then incubated with primary antibody anti-STAT3 (1:1,000, Cell Signaling, 30835) overnight at 4 °C. After washing 3x with TBST, membranes were incubated with the secondary anti-rabbit (IRDye800, 1:10,000, LI-COR, 925-32211). Blots were imaged on Bio-Rad ChemiDoc HMP Imaging System, and band densities were measured using NIH ImageJ FIJI software. STAT3 and p-STAT3 protein levels were normalized to GAPDH, which was used as a loading control. Activation of STAT3 was represented as the value of p-STAT3/STAT3 ratio for each sample. The mean ± SEM was calculated for each group.

### Immunofluorescence

#### Tissue Processing and Microtomy

Whole brains were harvested, bisected into two hemispheres, fixed overnight in 4% paraformaldehyde at 4°C, dehydrated in a series of graded TBS-Tween/ethanol solutions, and embedded in paraffin to obtain anterior-to-posterior coronal sections (n=1-2). Brains were trimmed to a depth of approximately 7000μm, or until the structures of the DG, CA1, CA2, CA3, and hilus could all be separately distinguished. 5μm coronal sections were collected. All sections for a given experiment (e.g., EZH2 induction in Saline WT, Saline EZH2nHom, KA WT, and KA EZH2nHom) were collected at comparable trim depths. All sections for a given experiment were collected on the same slide to reduce differences due to technical artifacts upon imaging. When this was not possible for experiments with many conditions (as in Fig. 9), a control section from the same saline-treated animal was included on every slide.

#### Immunostaining

Sections were subjected to immunofluorescence for NeuN, GFAP, total STAT3, IBA1, P2Y12, EZH2, and DAPI. Heat induced antigen retrieval was performed in a 10mM citric acid buffer (pH 6.0). After a combined permeabilization/blocking step, mouse anti-NeuN (1:200 ab104224 Abcam, or 1:500 MAB377 from Millipore), rabbit anti-STAT3 (1:500 #12640 Cell Signaling), rabbit anti-EZH2 (1:50 #5246 Cell Signaling), rabbit anti-IBA1 (1:600 #013-27691 Fujifilm), or rabbit anti-P2Y12 (1:200 #69766 Cell Signaling) primary antibodies were applied and incubated overnight at 4°C. Slides were washed with PBS then incubated with goat-anti-mouse Alexa Fluor 488 (ab150113 Abcam) and goat-anti-rabbit Alexa Fluor 594 (ab150080 Abcam) secondary antibodies at a concentration of 1:200 for one hour at room temperature, followed in some experiments by two-hour incubation at room temperature with Alexa Fluor 488-conjugated mouse anti-GFAP antibody (1:200 #53-9892-82 Invitrogen). Nuclear DNA was stained with a DAPI-containing mounting media (#H-1200 Vector Laboratories, Burlingame, CA).

#### Imaging

Representative images were captured at 10-20x magnification using a Nikon E600 fluorescent microscope fitted with an Olympus DP73 camera. DAPI was imaged with a V-2E filter with 340-380nm excitation range and 400nm long-pass/435-485nm band-pass emission filters. Proteins tagged with Alexa Fluor 488 were imaged with a B-2E/C filter with 465-495nm excitation range and 505nm long-pass/515-555 band-pass emission filters. Proteins tagged with Alexa Fluor 594 were imaged with a G-1B filter with 541-551nm excitation range and 565nm and 590nm long-pass emission filters. All images for a given marker were collected using identical settings and exposure times across conditions. Image capture was performed using Cellsens software (Olympus Life Science Solutions, Center Valley, PA).

#### Image Processing and Quantification

Image analysis and processing were performed in ImageJ (NIH, https://imagej.nih.gov/ij/). All images for a given marker were subjected to identical background subtractions, brightness/contrast (B/C) adjustments, thresholding, and particle analysis. Thresholds and settings were established with naïve sections and subsequently applied to all other groups. Grayscale TIFs from each channel were imported, and the corresponding RGB filter was applied. Background subtractions were performed, followed by brightness and contrast adjustments for figure panels.

Image processing for quantification involved background subtractions and the application of minimal B/C adjustments to generate single-channel images suitable for thresholding and the subsequent creation of binary images. Images were quantified by both intensity histograms and corrected total cell fluorescence (CTCF).

##### Intensity Histograms

The intensity histograms for each single-channel image were collected before thresholding, but after background subtraction and minimal B/C adjustments. Histograms quantified the frequency of pixels at a given intensity (0–255) for each image. Left-skewed histograms indicate a darker image, while right-skewed histograms indicate a brighter image. Histograms were exported and formatted to generate cumulative frequency distributions. Briefly, each intensity count was cumulatively summed such that each value in the series represented the total count up to that pixel intensity level (e.g., if counts for intensities 0, 1, and 2 were 1, 2, and 3, the cumulative values would be 1, 3, 6, and so on). The cumulative count for each intensity was then divided by the total pixel count (the final cumulative value for intensity 255) to normalize the distribution. This allowed for comparison of brightness levels, where images with left-skewed histograms would approach 100% at lower pixel intensities than images with right-skewed histograms. For those conditions (Saline, CP690550) with an n=2, cumulative counts were averaged. Results were graphed as cumulative frequency distributions in GraphPad Prism 9. The differences in these distributions were computed via Kolmogorov-Smirnov tests for analyses comparing two conditions or Kruskal-Wallis tests with Dunn’s correction for analyses comparing >2 conditions. These tests are standard for comparing cumulative distributions.

##### Corrected Total Cell Fluorescence (CTCF)

After background subtraction and minimal B/C adjustments, thresholding generated binary images, where regions of positive fluorescence were represented by white space and regions of no fluorescence by black space. The conditions for B/C adjustments and thresholding were set for saline groups first, and subsequently applied to all other conditions. Binary images were subjected to particle analysis to define regions of interest (ROIs). ROIs were called by ImageJ based on the binary image, rather than user input. ROIs were saved and subsequently applied to the processed images to quantify fluorescence in each ROI. Three identical ROIs were applied in areas of black space to quantify background fluorescence in the processed image. ROIs quantifying fluorescence outside of the hippocampus (e.g. see left-hand side of Fig. 9E *(XX)*) were omitted before measurement. Final measurements were exported for analysis. CTCF for each ROI was calculated and summed with the following formula:

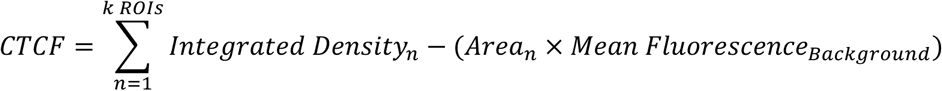

CTCF values for each condition were exported to GraphPad Prism 9 and depicted as bar graphs.

#### Animal care

All animal procedures and experiments were performed with approval from the University of Wisconsin-Madison School of Medicine and Public Health Instructional Animal Care and Use Committee, the Institutional Animal Care and Use Committee of Emory University, or the Institutional Animal Care and Use Committee of Tufts University, as appropriate, and according to NIH national guidelines and policies.

#### General

Male and female C57BL6 and FVB mice were bred and housed under a 12-hour light/dark cycle with access to food and water *ad libitum*. Mice were allowed to reach 4-6 weeks of age (C57BL6) or 5-7 weeks of age (FVB) before undergoing experimentation. Chemoconvulsant injections were performed at the same time of day (∼10 A.M. for kainate and 8-8:30A.M. for pilocarpine), and mice were returned to home cages before the start of the dark cycle (∼5 P.M.). Mice were housed with littermates, with males and females separated at weaning.

#### Transgenic mice

Cre-driver and EZH2 floxed mice were ordered from Jackson Laboratories (#003966, #022616, Bar Harbor, ME). The mice ordered were originally bred on a C57BL6 background. A second breeding colony was created in which mice were back-crossed for at least 10 generations into an FVB background. Details regarding the strain and sex used for every experiment can be found in Table S3. Using the previously described Synapsin1-Cre system(*35*) *EZH2^−/^*^−^; *Syn1Cre^+/−^* (EZH2nHom) mice were generated by mating *EZH2^flox/flox^; Syn1Cre^+/−^*females with *EZH2^flox/flox^* males (*125*). *EZH2^+/+^; Syn1Cre^+/−^* (EZH2nWT) mice were generated from mating pairs containing an *EZH2^flox/+^* male. To assess Cre expression, we used a reporter mouse with a lox-P flanked STOP cassette preventing transcription of tdTomato (#007909 Jackson Laboratories) (*126*). Reporter males were bred with *Syn1Cre^+/−^* females to generate Cre-positive animals and Cre-negative controls.

#### Seizure models

##### Repeated low-dose KA model

Mice were weighed and singly housed in observation chambers for the duration of the injections. Mice were injected intraperitoneally (i.p.) with synthetic kainic acid (KA) (7.5mg/kg for FVB, 5.0mg/kg for C57BL6) dissolved in 0.9% saline (#7065, Tocris Bioscience, Bristol, United Kingdom). At twenty-minute intervals, mice were given 7.5 or 5.0mg/kg injections of KA up to the third injection. The dosage was then reduced (5.0mg/kg for FVB, 3.75mg/kg for C57BL6). Animals continued to receive 5.0 or 3.75mg/kg injections of KA every twenty minutes until each reached SE. The subsequent KA injection was skipped if an animal experienced two or more Class V or VII seizures within a single twenty-minute cycle. Injections resumed for the next round, unless the animal reached SE. During induction of SE, seizures were scored using a modified Racine Scale (*127*) where I = freezing, behavioral arrest, staring spells; II = head nodding and facial twitches; III = forelimb clonus, whole-body jerks or twitches; IV = rearing; V = rearing and falling; VI = violent running or jumping behavior. Animals were considered to have reached SE after experiencing at least five Class V or VI seizures within a 90-minute window. KA mice were observed for 1-2 hours after SE was achieved. Animals were injected with 0.9% saline (s.c.), and soft gel food was provided in home cages. In the days following injection, animals were weighed and injected with 0.9% saline (s.c.) if body weight decreased by more than 0.5g.

##### Pilocarpine model of status epilepticus

Status epilepticus was induced in male and female C57BL/6CRN mice with pilocarpine (8-12 weeks old) (*128, 129*). Mice were injected with terbutaline (4 mg/kg i.p., Sigma T2528) and methylscopolamine (4 mg/kg ip, Sigma S8502) to alleviate respiratory and cardiovascular effects of pilocarpine, then 30 min later received saline or pilocarpine (280 mg/kg i.p. as free base, Sigma P6503) as described (*32*). After 1 hour of SE, seizures were interrupted by diazepam (10mg/kg i.p.). Mice were weighed and scored for behavioral deficits (Irwin score) daily, with injections of lactated Ringer’s solution as needed.

##### IHKA model of status epilepticus

The intrahippocampal kainic acid (IHKA) model was employed to generate chronically epileptic mice as previously described (*130*). Kainic acid (100nL of 20 mM; vIHKA) or vehicle (100 nL of sterile saline; vIHSA) was stereotaxically injected into the hippocampus using the following coordinate (from Bregma): 3.6 mm posterior, −2.8 mm lateral, and 2.8 mm depth. Only mice which experienced status epilepticus and went on to develop spontaneous recurrent seizures were used in the current study.

#### Fluorothyl and PTZ seizure threshold testing

Seizure threshold tests were performed at the same time of day (∼10 A.M.), and mice were returned to home cages before 5 P.M. All seizure threshold tests were video recorded, and seizure behavior was scored by a blinded observer. Flurothyl seizure threshold tests were performed on male and female EZH2nWT and/or EZH2nHom mice using 100% bis(2,2,2-trifluoroethyl) ether (#287571, Sigma-Aldrich, St. Louis, MO). All tests were performed in a fume hood. Mice were placed in an airtight 10L. Plexiglas chamber (8.5”X10.75”X6.75”). Fluorothyl was infused into the chamber via a peristaltic pump at a rate of 40μL/minute onto a piece of Whatman filter paper placed at the top of the chamber. The time to generalized tonic clonic seizure (GTCS) was recorded from the start of fluorothyl infusion. Mice reached GTCS when they exhibited a complete loss of postural/motor control. After experiencing a generalized tonic clonic seizure, mice were rapidly removed from the chamber, and fluorothyl infusion was terminated. Mice were transferred to a recovery cage and kept isolated until motor control and normal behavior returned (∼10 minutes). Mice were returned to home cages after recovery. For PTZ threshold tests, PTZ was dissolved at 5mg/mL in 0.9% normal saline, and a single 45mg/kg dose was administered (i.p.). Time to generalized tonic clonic seizure was recorded.

For Fig. S11, flurothyl or PTZ seizure threshold tests were performed once on naïve mice pre-treated with vehicle (i.p.) or CP690550 (15mg/kg i.p.) 30 minutes prior to testing. For Fig. S1, two flurothyl seizure threshold tests per mouse were performed: the first at one week prior to kainate injections and the second at four days post-SE.

#### CP690550 citrate treatment of KA mice

Following status epilepticus, an independent researcher (not the experimenter) randomly assigned mice to drug and vehicle conditions, prepared both solutions, and aliquoted one or the other into tubes labeled with the mouse ID only. The independent researcher did not participate in any seizure threshold testing or spontaneous seizure video scoring. CP690550 citrate (tofacitinib citrate) (#4556, Tocris Bioscience, Bristol, United Kingdom) was dissolved in a solution of 70% saline (0.9%)/15% ethanol/15% DMSO. The 70% saline/15% ethanol/15% DMSO solution was administered as vehicle. In acute experiments, CP690550 or vehicle was delivered to mice (i.p.) at 6, 18, 24, 48, and 72 hours post-SE. The first dose of drug was given at 25mg/kg; all others were 15mg/kg. In chronic experiments, CP690550 or vehicle was delivered daily for two weeks at 15mg/kg (i.p.).

#### Spontaneous Recurrent Seizure (SRS) Detection

##### Long-term video monitoring for spontaneous seizures

Male and female FVB mice (WT, EZH2nWT, and EZH2nHom) were singly housed in observation chambers for the duration of recording with access to food and water. During CP690550 treatment, mice were weighed daily and injected before recording began. Mice that did not exhibit at least 1 seizure per week (0.2/day) during the baseline recording period or who died during recording were triaged from the analysis. Deaths during the chronic period after vehicle or CP690550 treatment were noted and analyzed for any treatment effects (Fig. S12).

Videos were reviewed and scored by an experimenter blinded to treatment and genotype using the modified Racine scale. In the experiments described by Figure 6, only Racine class IV-VI seizures were recorded. For each mouse, daily seizure burden was calculated as:

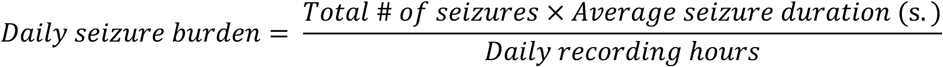

Mice that did not respond to CP690550 treatment with at least a 50% reduction in seizure frequency and burden were classified as non-responders and omitted from the disease modification phase of the experiment. SRS analyses including non-responders can be found in Fig. S12, and Table S1 includes mean seizure frequency, burden, and metadata for each animal. Of the 51 mice described in the seizure analysis, five CP690550-treated mice were non-responders and therefore omitted from the disease modification arm of the experiment. Random harvests on the final day of treatment decreased sample sizes from n=26 to n=20 for vehicle-treated mice, and from n=25 to n=18 for CP690550-treated mice. Random harvests two weeks post-washout further decreased sample sizes from n=20 to n=15 for vehicle-treated mice, and from n=18 to n=12 for CP690550-treated mice.

##### Continuous EEG recording and seizure detection

At the time of stereotaxic IHKA or IHSA injections, mice were implanted with chronic EEG recording headmounts (Pinnacle Technology, cat. #8201) which were affixed to the skull with 4 screws: 2 screws serving as EEG leads placed in the anterior and posterior cortex, 1 as a reference ground, and 1 as the animal ground. The headmounts were secured with dental cement. 24/7 EEG recordings were collected at 4 KHz using a 100X gain preamplifier high pass filtered at 1KHz (Pinnacle Technology, cat. #8292-SE) and tethered turnkey system (Pinnacle Technology, cat. #8200).

Seizures were detected using a custom analysis pipeline developed in-house in python (seizy; https://github.com/neurosimata/seizy) as previously described (*130*). Briefly, EEG traces were automatically detected using a thresholding method from traces that were downsampled (decimated) to 100 Hz and divided into 5 second segments. Power (2-40 Hz) and line length were used to identify seizures. Seizures were only included if two consecutive 5 second segments were classified as seizures (minimum 10 seconds length) and were manually verified by an experienced investigator with expertise in electrographic seizure detection. Seizure frequency was calculated as the # of seizures per day by measuring total number of seizures a mouse had over a 24-hour period. The average seizure duration was calculated by taking the mean seizure duration for each mouse in each recording period. For each mouse, seizure burden was calculated as:

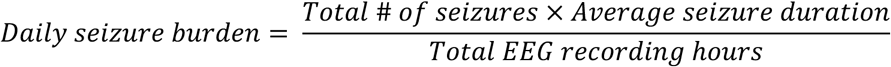

#### Y-maze cognitive testing

Working memory spontaneous alternation (SA) and short-term memory forced alternation (FA) tests were performed at 3 time points: one (FA) or two (SA) weeks prior to induction of SE, post-CP690550 treatment (12 weeks post-SE), and post-washout (14 weeks post-SE). All tests were video recorded and analyzed by an experimenter blinded to treatment. Mice were habituated in the testing room for 30 minutes prior to all trials. We used spontaneous alternation to evaluate working memory and forced alternation to evaluate short-term memory; both tests occurred within a three-arm Y-maze.

##### Spontaneous Alternation

For spontaneous alternation, mice were placed in the starting arm of the maze and allowed to explore freely for 8 minutes. Sequential arm choices during exploration of a three-arm Y-maze are considered an indicator of spatial working memory because the innate curiosity of a healthy mouse promotes exploration of the arm least recently visited (*32, 53, 54*). The number of choices resulting in a spontaneous alternation (e.g., an entry sequence of ABC but not ACA) was expressed as a percentage of the total number of choices. After prolonged time in the Y-maze, habituation to novelty, and thus changes in motivation, can reduce the percentage of spontaneous alternations over time (*32*). Testing naïve mice, we observed the transition from an exploratory phase to an escape phase, and the total number of alternations for each mouse ranged from 20-100. To mitigate the effects of habituation and the variance in total alternation, we analyzed only the first 15 alternations of each mouse. Mice that did not reach 15 alternations were removed from the dataset. Distance traveled during each 8-minute test was measured to ensure that deficits in SA post-SE were not due to mobility impediments.

##### Forced Alternation

Forced alternation (FA) tests depend on a mouse’s ability to recall and apply spatial short-term memory, motivated by their innate curiosity. FA tests involved two 5-minute trials 30 minutes apart: a sample trial and a retrieval trial. In the sample trial, the mouse was placed in the starting arm and allowed to explore the maze with one arm blocked off. Between trials the maze was washed with 70% EtOH. In the retrieval trial, the previously blocked arm was opened, and the mouse was allowed to explore freely for 5 minutes. Mice that did not enter all three arms of the maze during the retrieval trial of FA tests were excluded from the dataset. FA tests were quantified as percentage of time spent in the novel arm. A mouse was considered inside the novel arm when its hindlimbs entered it, and outside the novel arm when its hindlimbs exited. Time spent in the center of the maze was not counted.

### Statistical analysis

For all statistical analysis – unless otherwise specified – q<0.05 or p<0.05 corrected for multiple comparisons was considered statistically significant (*p/q<0.05, **p/q<0.01, ***p/q<10^−3^, ****p/q<10^−4^). Data were graphed as mean ± standard deviation (SD) in Prism 9 software (Graphpad Software, La Jolla, CA). Statistical tests were also performed in Prism. Western blot, seizure threshold, and spontaneous seizure data was analyzed by t-test, 2-way ANOVA/Mixed effects models, or 1-way ANOVA, with Tukey’s correction for multiple comparisons or the two-stage step-up method of Benjamini, Krieger, and Yekutieli (abbreviated BKY in figure legends). Paired tests were used where appropriate. Cumulative frequency distributions were measured by Kolmogorov-Smirnov tests for unpaired data with 2 conditions, Wilcoxon matched pairs signed rank test for paired data with 2 conditions, and Krusak-Wallis tests with Dunn’s correction for data with >2 conditions. All statistical tests were two-tailed.

## Supporting information

SRS_data

mouse_human_clusters_comparisons

mouse_metadata

## Acknowledgements

We thank Tristan Holz for 3D printing the Y-maze.

## Supplemental Figures

**Fig. S1.**
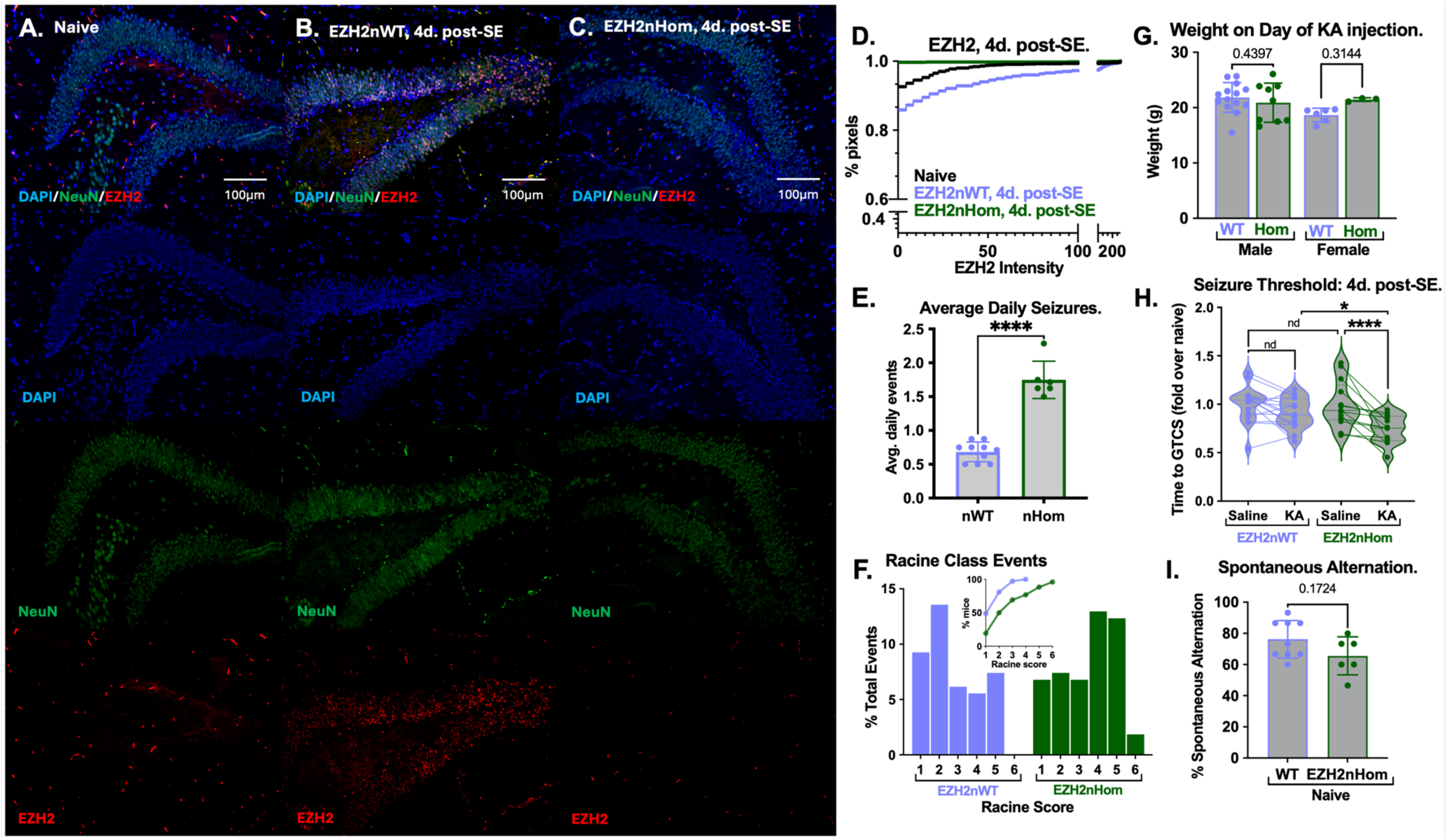
Neuronal EZH2 induction after status epilepticus mitigates chronic disease progression in the systemic kainate model. **(A-C)** Merged and single-channel immunofluorescence images of the dentate gyrus labeling DAPI, NeuN, and EZH2 on day 4 post-SE and **(D)** quantitation of EZH2 histograms from each panel. Statistical analysis was performed by Kruskal-Wallis test with Dunn’s correction for multiple comparisons. Naïve mice showed low levels of EZH2 in neurons, while wildtype mice injected with kainate and harvested on day 4 post-SE exhibited neuronal induction of EZH2 (p<10^−4^). EZH2 homozygous neuronal knockout mice failed to show this induction, with EZH2 levels reduced compared to EZH2nWT mice 4d. post-SE (p<10^−4^). **(E)** SRS activity was video recorded in epileptic EZH2nWT and nHom mice 5-7 weeks post-SE. EZH2nHom mice showed a 2.5-fold increase in seizure frequency (unpaired t-test; n=6-10). **(F)** Racine scale events for EZH2nHom and EZH2nWT mice were scored and quantified as % total events (inset: cumulative frequency distribution). EZH2nHom mice had more severe seizures than WT mice. (Kolmogorov-Smirnov test; D:0.833; n=6-10). **(G)** Neuronal EZH2 deletion did not alter weight in male or female mice. **(H)** Flurothyl seizure threshold was measured in EZH2nWT and nHom mice 1 week before systemic KA injection and again 4 days post-SE. Deletion of an EZH2 allele does not alter naïve seizure threshold (q>0.05, 2-way ANOVA with BKY correction) (n=13-16). While EZH2nWT mice did not yet show reduced seizure threshold on day 4 post-SE (9% reduction, q>0.05), EZH2nHom mice showed a 26% reduction (q<10^−4^), a 16% reduction compared to WT mice (q<0.05). **(I)** Working spatial memory was measured by spontaneous alternation testing one week before KA injection. Neuronal deletion of EZH2 did not alter working memory. *(**Seizure Model:** Systemic kainate; **Seizure detection:** Video recording)*

**Fig. S2.**
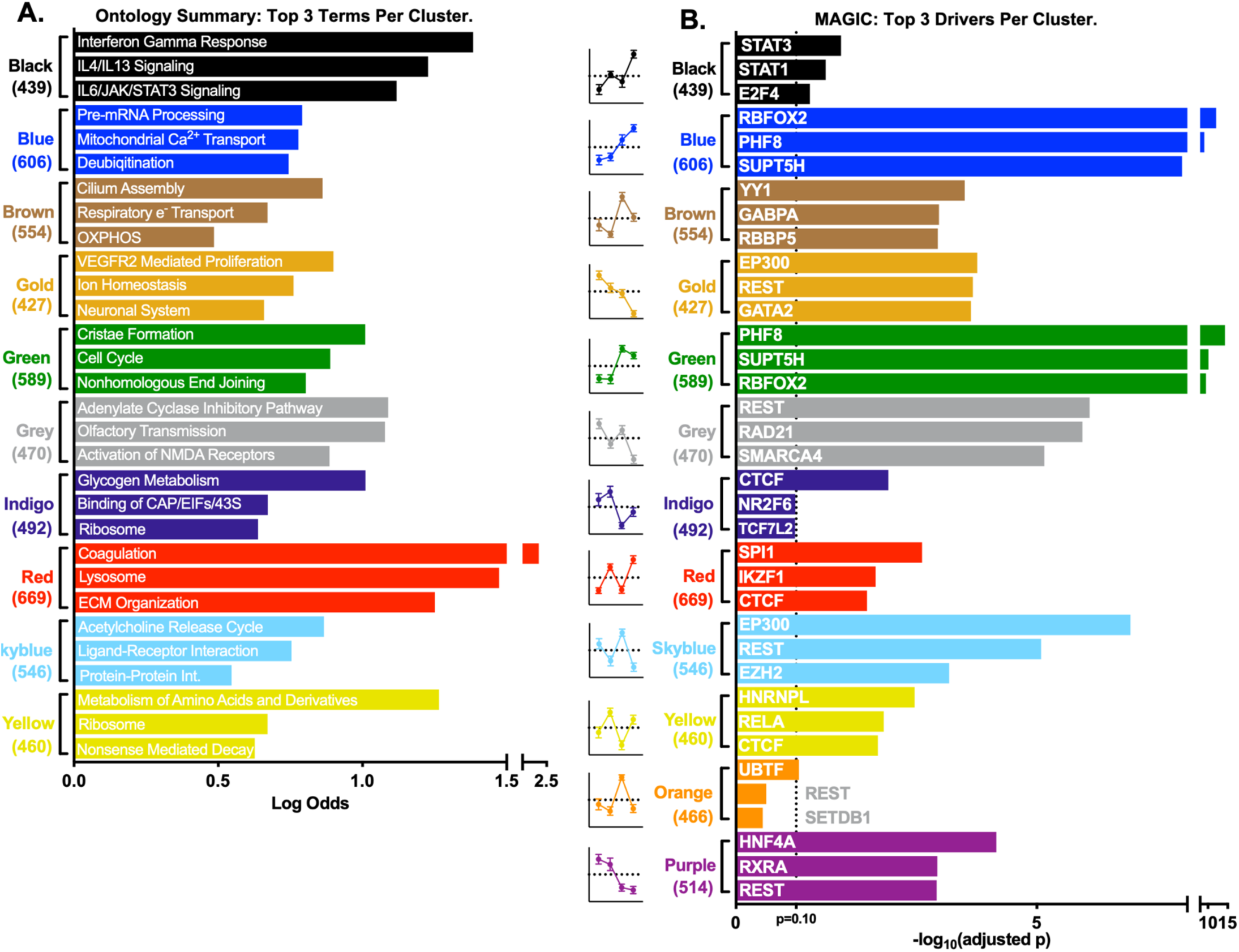
Ontology and MAGIC summaries for DEGs across all Leiden clusters generated by bulk RNAseq. Leiden clusters from 6241 differentially expressed genes (fold change>1.2x, adjusted p<0.01) across saline, kainate, EZH2nWT, and EZH2nHom conditions were subjected to **(A)** ontological and **(B)** MAGIC analysis. Genes upregulated after SE were housed in the Black, Blue, Indigo, Red, and Yellow clusters, with ontological profiles generally related to inflammation, translation, post-translational modification, and glycogen metabolism. Genes downregulated after SE were housed in the Brown, Gold, Green, Grey, Skyblue, Orange, and Purple clusters, with ontological profiles generally related to cell proliferation, RNA processing, neurotransmitter release, and excitatory neurotransmission. The Orange and Purple cluster genes had no hits across the KEGG, Reactome, Biocarta, or Hallmark databases. For the upregulated clusters (Black, Blue, Indigo, Red, Yellow), the top factors predicted to drive gene expression for each cluster were STAT3, RBFOX2, CTCF, SPI1, and HNRNPL, respectively. For the downregulated clusters (Brown, Gold, Green, Grey, Skyblue, Orange, Purple), the top factors predicted to drive gene expression for each cluster were YY1, EP300, PHF8, REST, EP300, UBTF, and HNF4A, respectively. *(**Seizure Model:** Systemic kainate)*

**Fig. S3.**
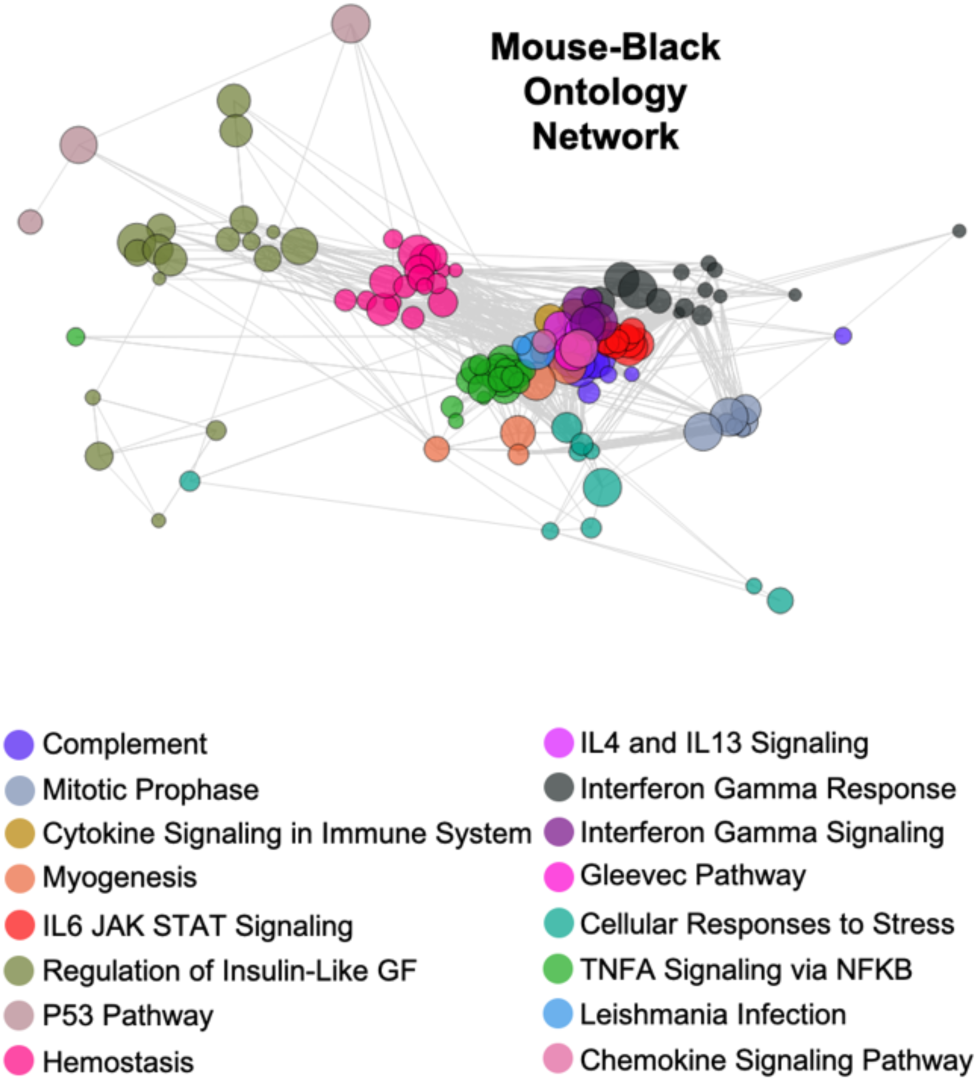
Ontology network example for the mouse-Black cluster. The 439 mouse-Black cluster genes were compared to gene lists associated with terms in KEGG, Reactome, Biocarta, and Hallmark databases; terms with an odds ratio>1 and p<0.05 across all databases were sorted into ten communities via a network approach. Community names are the ontological term with the highest odds ratio in the community. (***Seizure Model:*** *Systemic kainate)*

**Fig S4.**
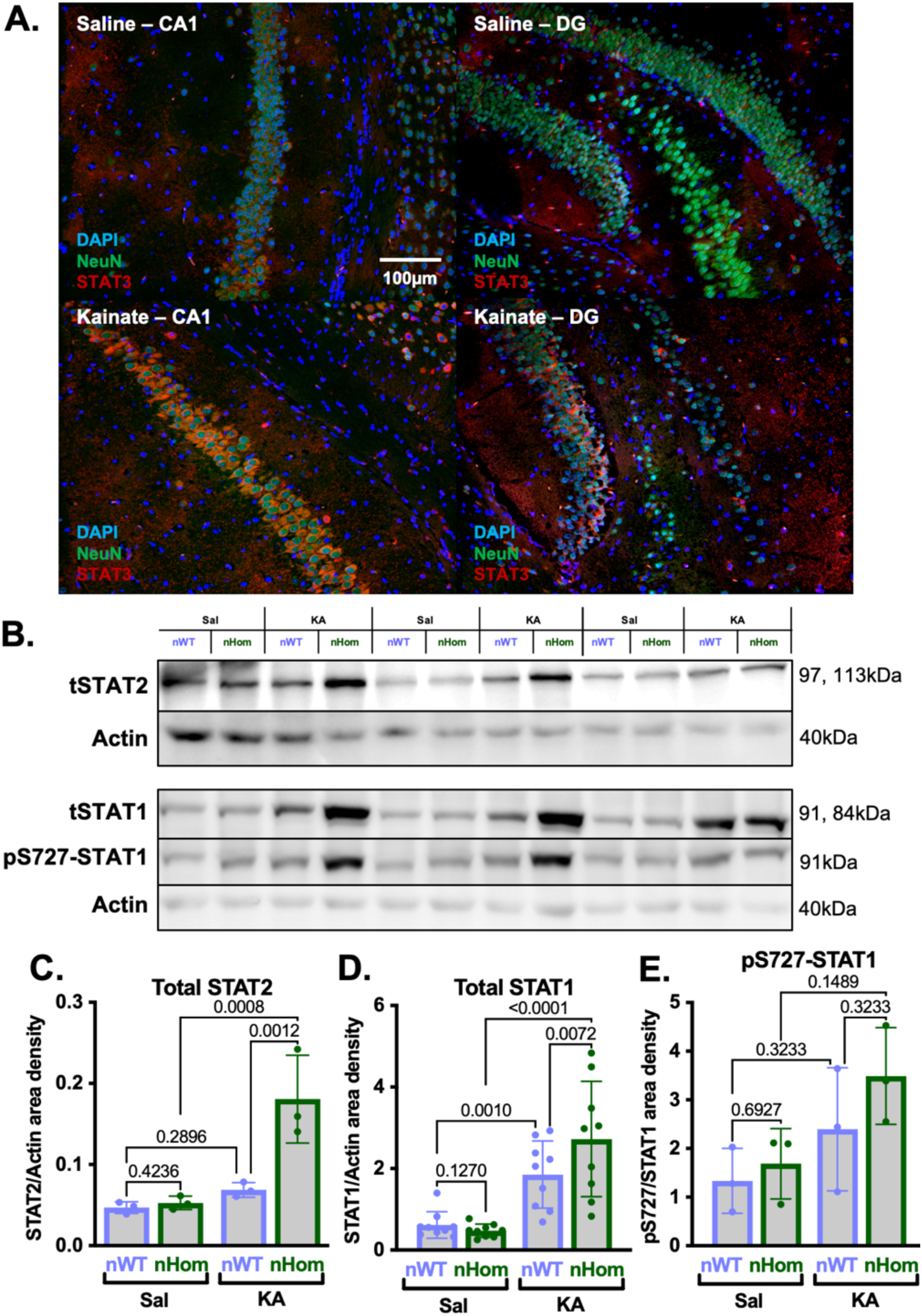
STAT3 is induced in neurons, and other STAT family proteins are hyper-induced in EZH2nKO mice. **(A)** 20x images of the CA1 pyramidal cell layer (CA1) and dentate gyrus (DG). Hippocampal slices were collected on day 1 post-SE from saline- and KA-injected mice; immunofluorescence was performed to detect nuclear DNA, NeuN, and total STAT3. **(B)** Representative western blots for total STAT2, total STAT1, phospho-STAT1 (Ser727) and beta-actin on day 4 post-SE. For all proteins quantified in panels B-D, deletion of an EZH2 allele did not alter protein levels in naïve animals. **(C)** Total STAT2 was induced after SE in EZH2nHom mice by 3.4-fold, or 2.6-fold greater than EZH2-sufficient littermates (nWTs) post-SE (2-way ANOVA with Tukey’s correction; n=3). **(D)** Total STAT1 protein was induced by 5.7-fold in EZH2nHom mice post-SE, or 1.5-fold greater than WTs post-SE (2-way ANOVA with Tukey’s correction; n=9). **(E)** pS727-STAT1 was not robustly altered across the four conditions (2-way ANOVA; n=3). *(**Seizure Model:** Systemic kainate)*

**Fig. S5.**
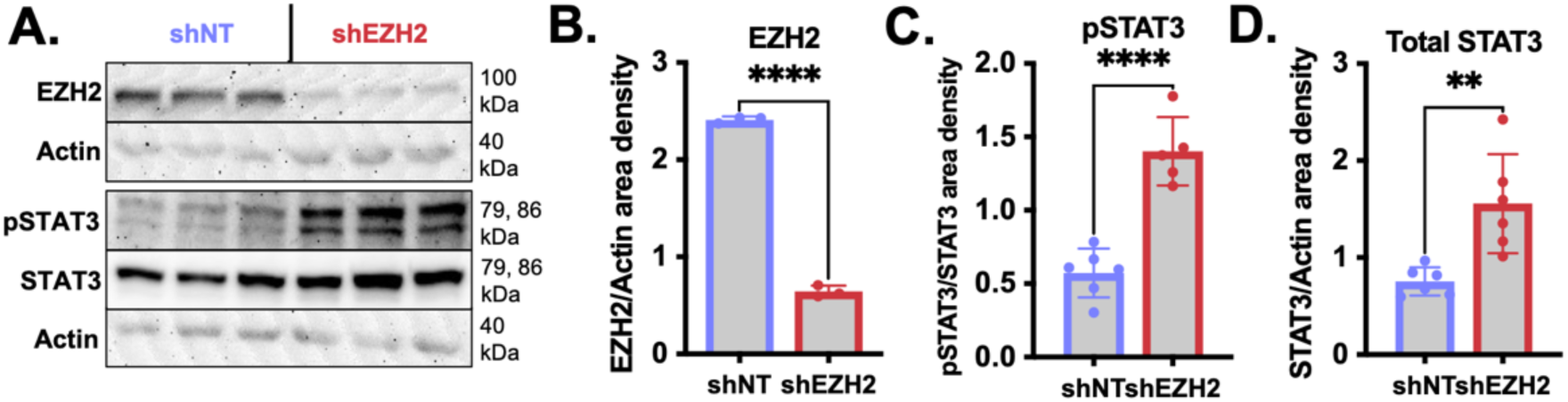
EZH2 is required to temper JAK/STAT signaling N2A cells. **(A)** N2A cells were stably transfected with EZH2 shRNA (shEZH2) or non-targeting shRNA (shNT). Protein was harvested from shNT and shEZH2 cells for western blot analysis. **(B)** EZH2 was reduced 3.7-fold in the knockdown cells (unpaired t-test; n=3). We observed **(C)** a 2.4-fold induction of pSTAT3 (unpaired t-test; n=5-6) and **(D)** a 2.1-fold induction of total STAT3 in shEZH2 cells (unpaired t-test; n=6).

**Fig. S6.**
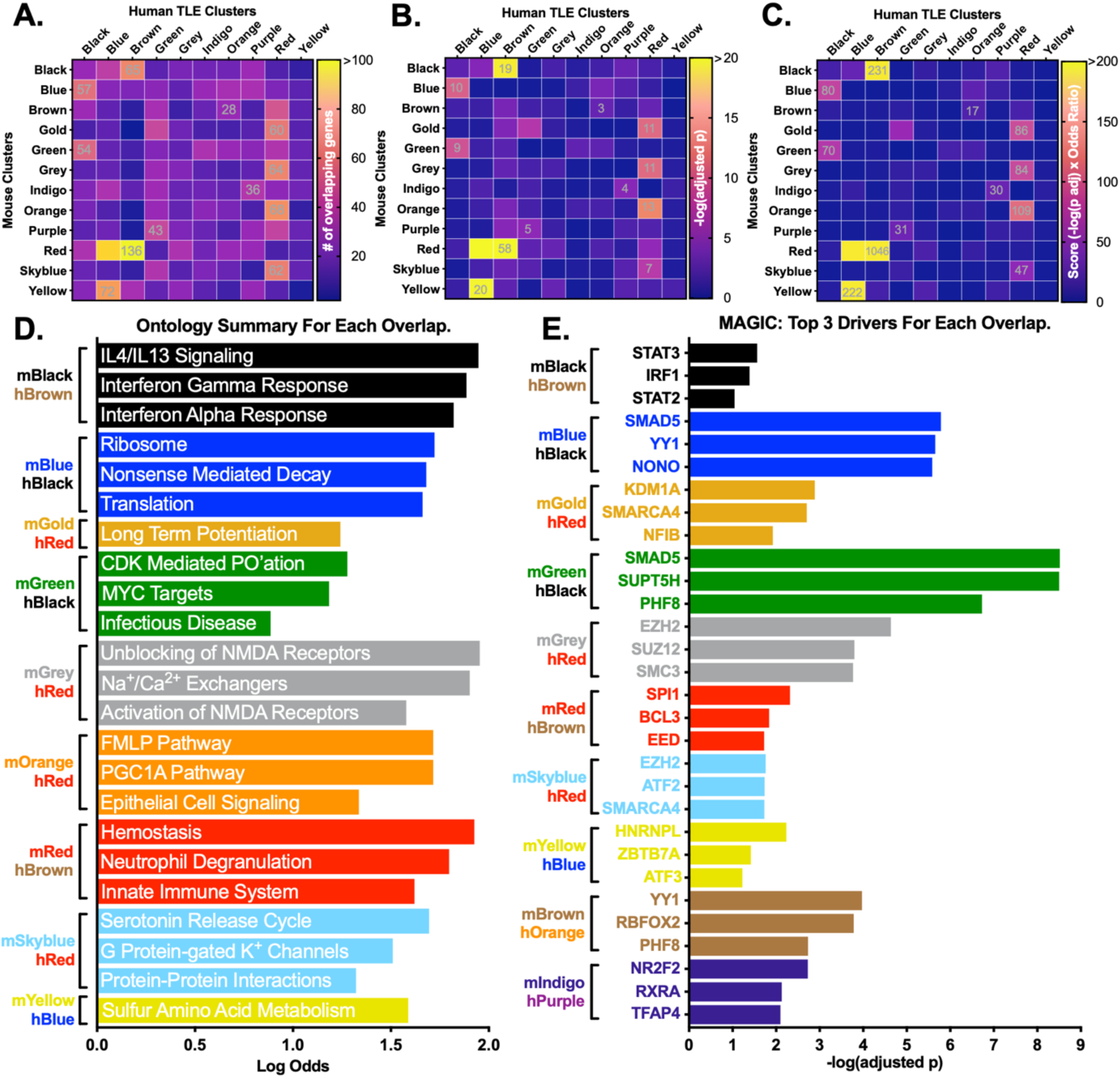
Ontology and MAGIC summaries for gene overlaps across all mouse clusters and human TLE clusters. Leiden clusters from saline, kainate, EZH2nWT, and EZH2nHom mice were subjected to Fisher exact tests with the Benjamini Hochberg correction for multiple comparisons, quantifying the overlap of each mouse cluster with 10 clusters of genes expressed across human TLE by **(A)** number of overlapping genes, **(B)** confidence (-log(adjusted p)), and **(C)** score (confidence * Odds ratio). The human cluster with the greatest score for each mouse cluster was selected for downstream **(D)** ontological and **(E)** MAGIC analysis. *(**Seizure Model:** Systemic kainate)*

**Fig. S7.**
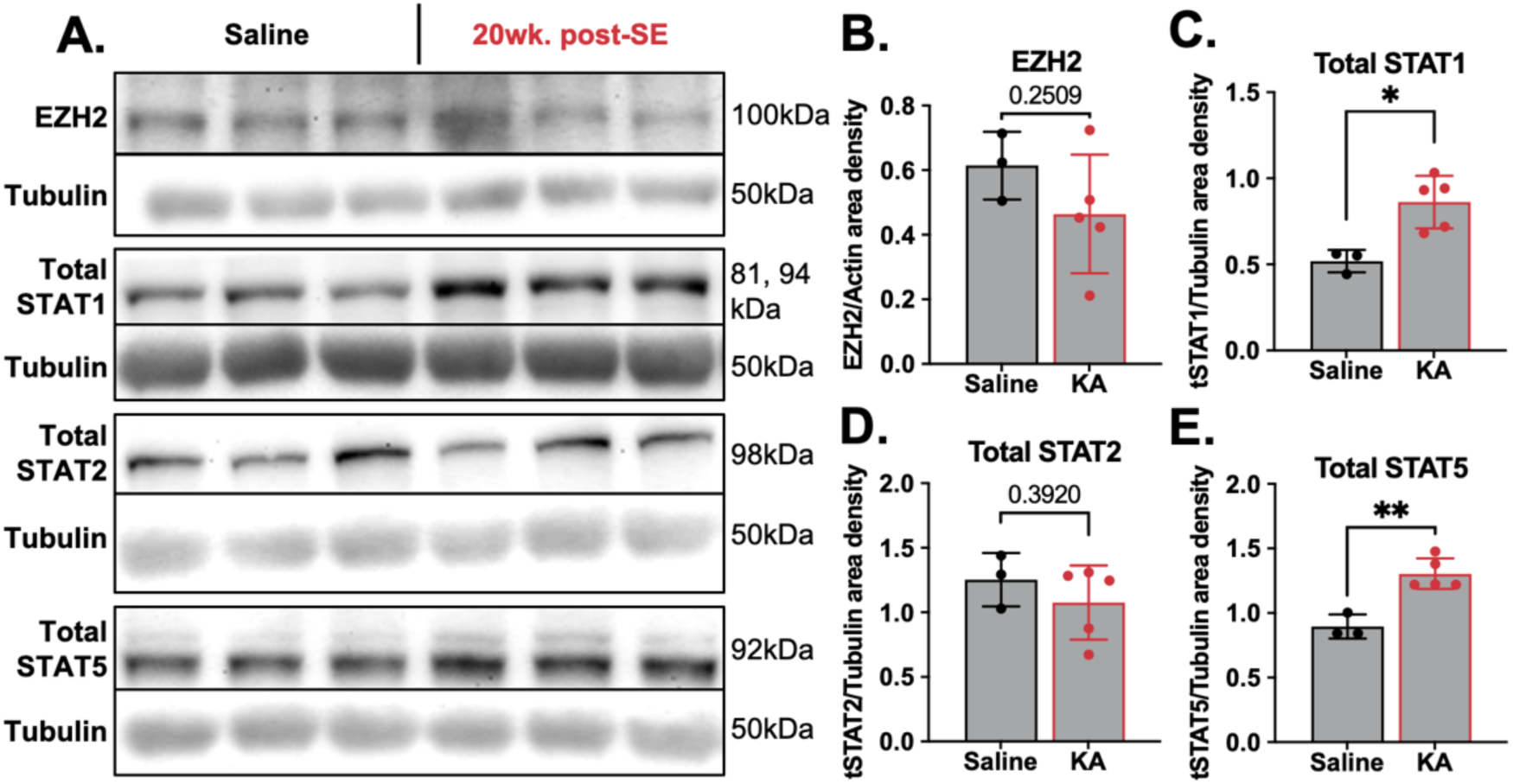
Protein expression of JAK/STAT pathway components in chronic epilepsy. **(A)** Hippocampi were harvested from 20wk. post-SE chronically epileptic mice and naïve controls. Protein lysates were subjected to western blotting. **(B)** EZH2 was unchanged from naïve levels in chronic epilepsy (unpaired t-test; n=3-5). At 20 weeks post-SE compared to naïve mice, **(C)** Total STAT1 was induced 1.7-fold (p<0.05, n=3-5), **(D)** total STAT2 was unchanged (n=3-5), and **(E)** total STAT5 was induced 1.5-fold (p<0.01, n=3-5). *(**Seizure Model:** Systemic kainate)*

**Fig. S8.**
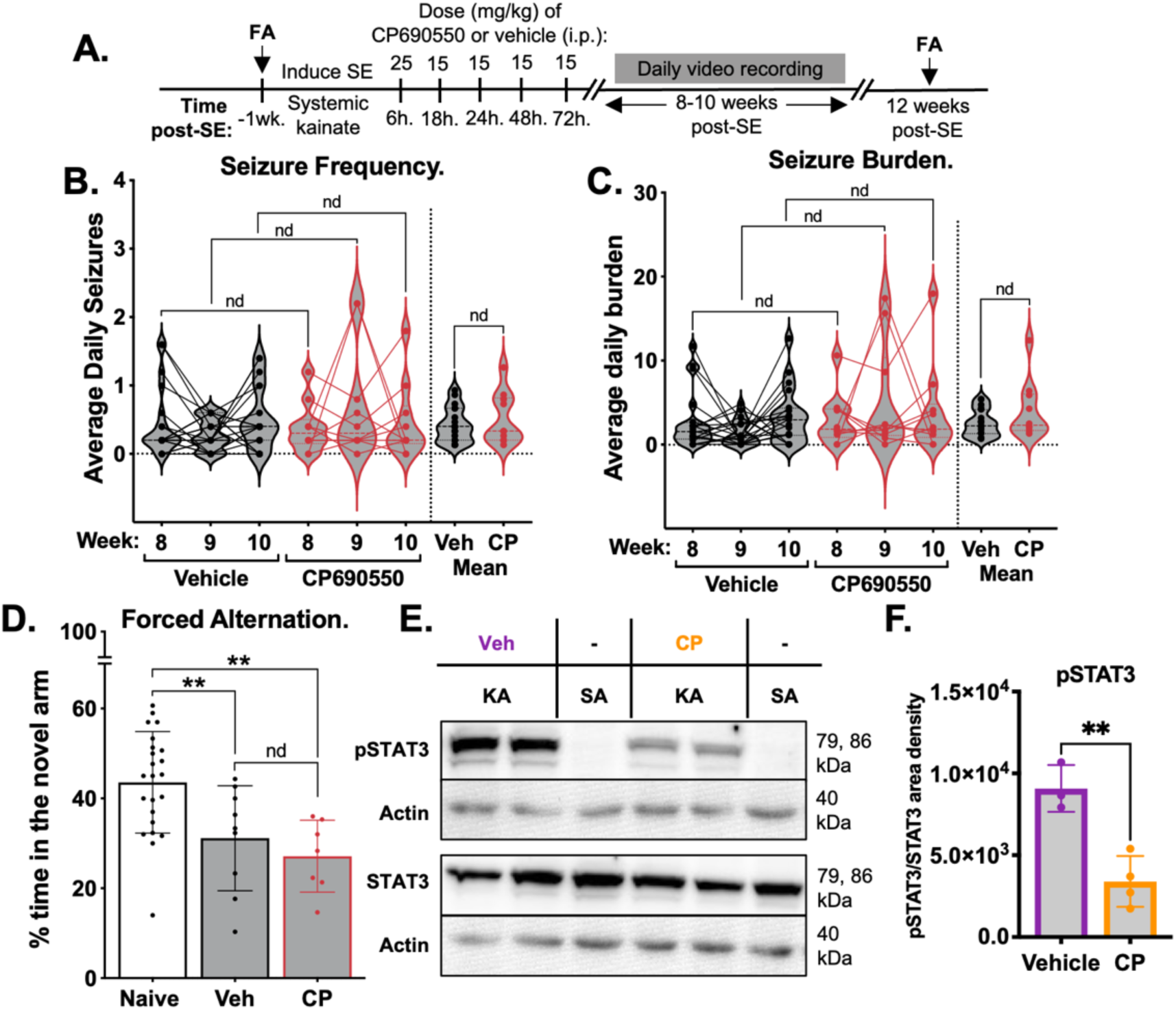
Transient CP690550 treatment acutely post-SE does not alter epileptogenesis in wildtype mice. **(A)** SE was induced via <u>systemic kainate</u>, and wildtype mice were treated with five doses of either vehicle or CP690550 up to 72h. post-SE. Behavioral spontaneous seizure activity was monitored by <u>video recording</u> from weeks 8-10 post-SE. Seizure frequency **(B)** and burden scores **(C)** across vehicle- and CP690550-treated mice were quantified. Acute treatment with CP690550 was not antiepileptogenic (2-way ANOVA with BKY correction; n=10-20). No week-to-week differences in seizure frequency or burden were detected in vehicle-versus CP690550-treated mice, nor in the average seizure frequency or burden across weeks. **(D)** The same mice were subjected to forced alternation Y-maze testing 1 week before SE and 12 weeks post-SE. Acute treatment with CP690550 did not rescue epilepsy-associated short term memory decline (1-way ANOVA with BKY correction; n=7-23). Chronically epileptic mice showed reduced performance in the forced alternation test regardless of treatment acutely post-SE (**Vehicle:** 29% reduction; **CP690550:** 38% reduction). **(E)** After induction of SE and three doses of CP at 6, 12, and 24h. post-SE (25, 15, and 15mg/kg i.p., respectively), mice were sacrificed 3 hours after the last injection to confirm drug target engagement via western blot. Naïve mice (lanes 3,6) were included to confirm pSTAT3 induction typically observed on day 1 post-SE. **(F)** Drug-treated animals exhibited a 2.7-fold reduction in pSTAT3 compared to vehicle-treated animals (unpaired t-test; n=3). *(**Seizure Model:** Systemic kainate; **Seizure detection:** Video recording)*

**Fig. S9.**
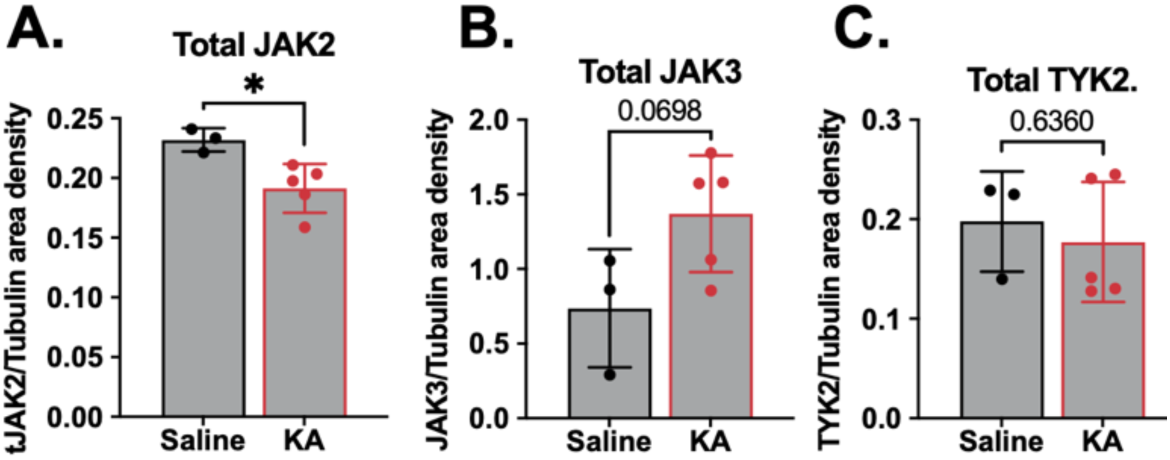
Western blot quantitations for JAK2, JAK3, and TYK2 in chronic epilepsy. Hippocampi were harvested from 20wk. post-SE chronically epileptic mice and naïve controls. Protein lysates were subjected to western blotting. **(A)** Total JAK2 expression was reduced 1.2-fold in chronically epileptic mice from naïve levels (p<0.05, n=3-5). **(B-C)** Total JAK3 and total TYK2 were unchanged, although total JAK3 trended towards induction (n=3-5). *(**Seizure Model:** Systemic kainate)*

**Fig. S10.**
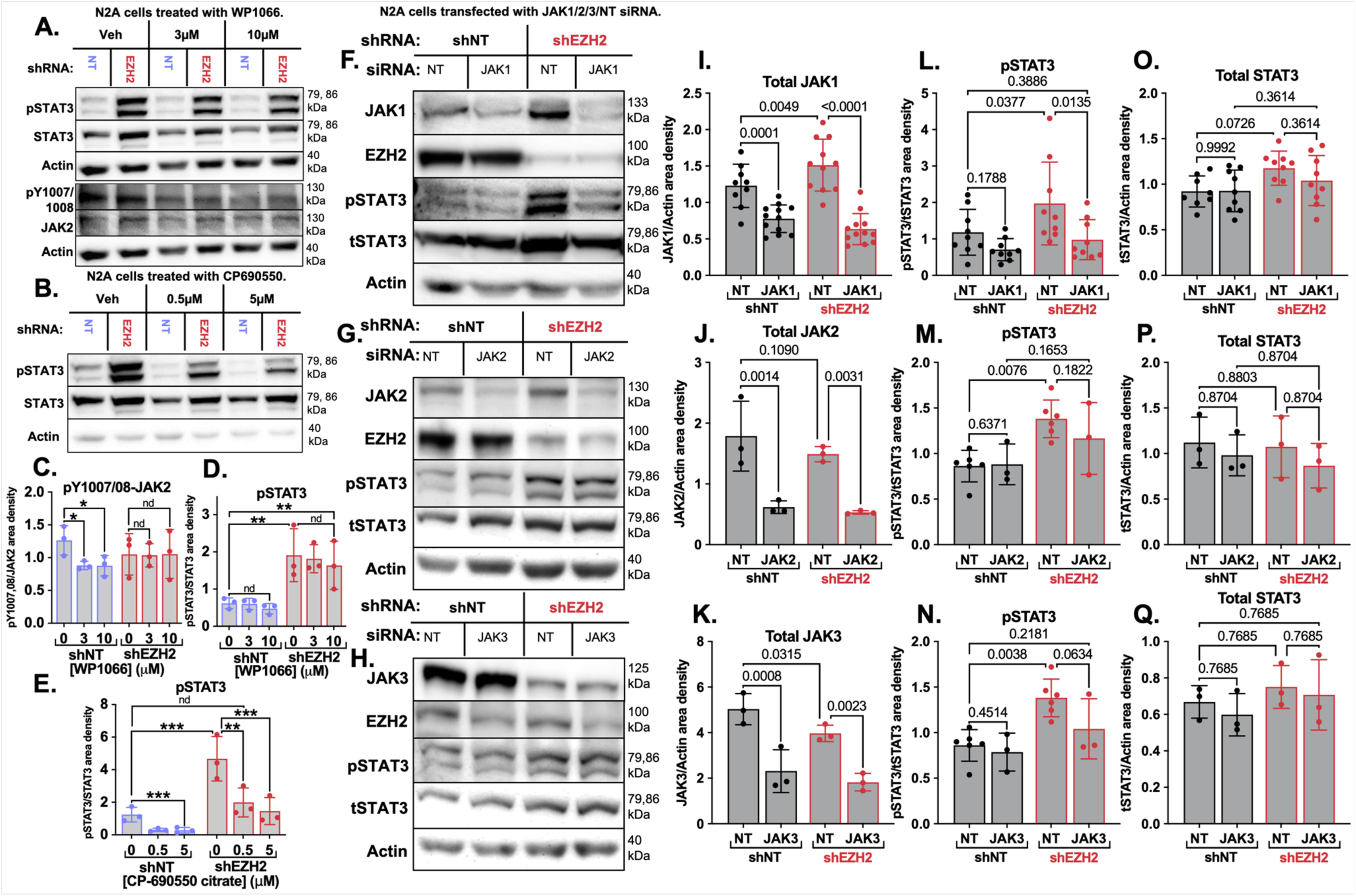
JAK1 is required for hyper-activation of STAT3 in the absence of EZH2. **(A)** shNT and shEZH2 cells were treated with the JAK2/STAT3 inhibitor WP1066 to a final concentration of 0 (DMSO), 3, or 10μM for one hour. **(B)** shNT and shEZH2 cells were treated with the JAK1/3 inhibitor CP690550 citrate to a final concentration of 0 (DMSO), 0.5, or 5μM for one hour. **(C)** WP1066 reduced pY1007/08-JAK2 in shNT cells for both concentrations. No change was observed in knockdown cells. **(D)** pSTAT3 was induced in vehicle-treated shEZH2 cells by 3-fold (q<0.01) in the WP1066 experiment. WP1066 did not alter pSTAT3 levels in control or knockdown cells at any concentration. **(E)** pSTAT3 was induced in vehicle-treated shEZH2 cells by 3.7-fold (q<10^−3^) in the CP690550 experiment. CP690550 treatment for one hour reduced pSTAT3 levels of KD cells by 2.3-fold at 0.5μM (q<0.01) and 3.2-fold at 5μM (q<10^−3^). At 0.5μM CP, pSTAT3 levels in KD cells were reduced to that of vehicle-treated shNT cells (2-way ANOVA with BKY correction; n=3). In a parallel experiment, protein was harvested for western blot analysis from shNT and shEZH2 cells transfected with either non-targeting or JAK1 **(F),** JAK2 **(G)**, or JAK3 **(H)** siRNAs, resulting in four conditions (shNT:siNT, shNT:siJAKx, shEZH2:siNT, shEZH2:siJAKx). Results for panels I-Q were analyzed by 2-way ANOVA with BKY’s correction (n=3-12). Both shNT and shEZH2 cells showed reduction of JAK1 **(I)**, JAK2 **(J)**, and JAK3 **(K)** (compare bars 1 and 2, 3 and 4, for each panel). **(I)** Only JAK1 was induced with EZH2 knockdown (1.2-fold, q<0.01) (compare bars 1 and 3). **(L-N)** All experiments showed induction of pSTAT3 with EZH2 knockdown (compare bars 1 and 3 for each panel). **(L)** Only JAK1 RNAi succeeded in reducing hyperactivated pSTAT3 (2-fold, q<0.05) to levels indistinguishable to that of shNT:siNT cells (p>0.05, compare bars 3 and 4, 1 and 4). **(O-Q)** Total STAT3 was not altered across any conditions in this experiment.

**Fig. S11.**
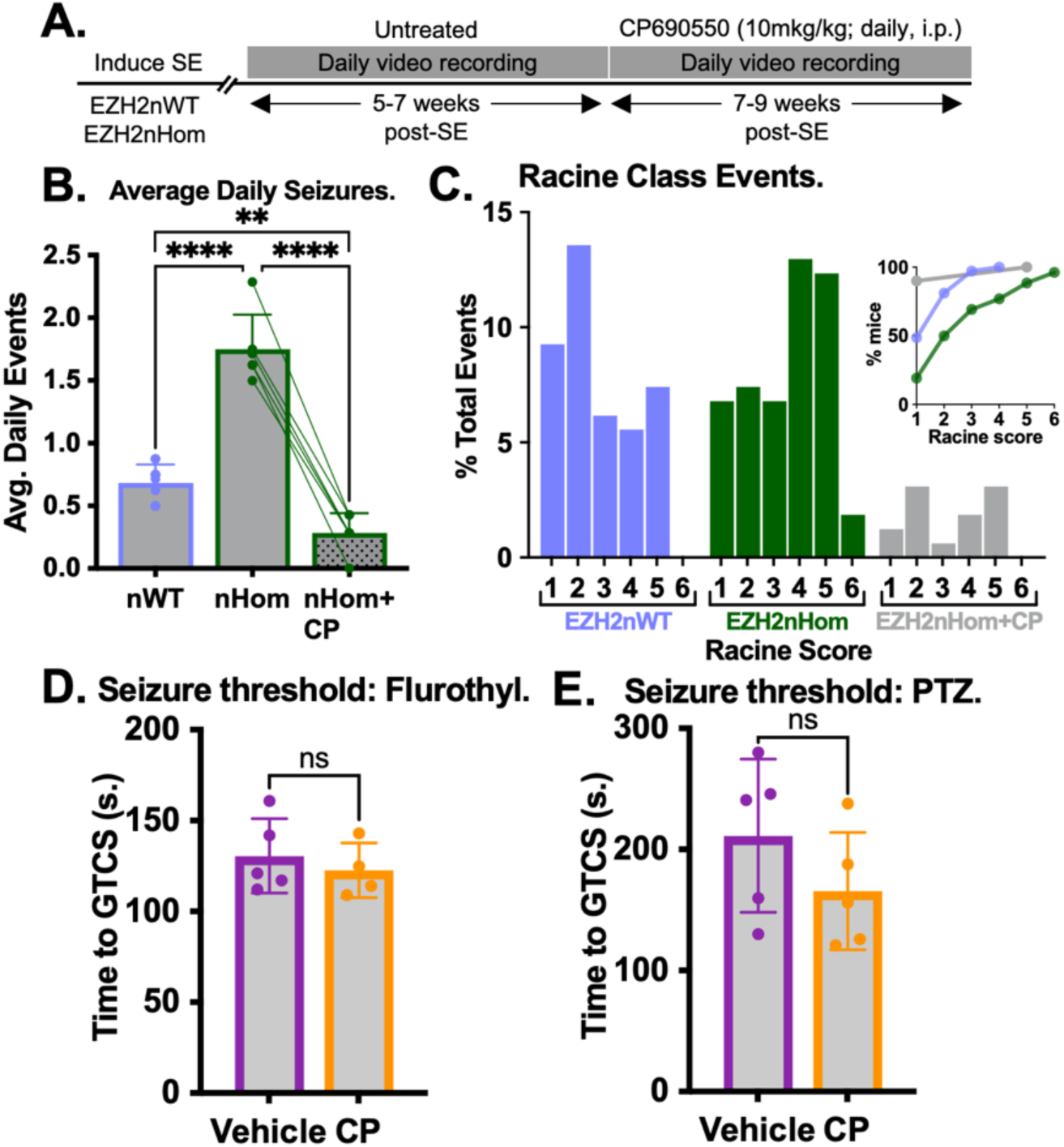
CP690550 treatment of chronically epileptic EZH2nHom mice rescues exacerbated epileptic phenotypes. **(A)** After baseline SRS activity was monitored from 5-7 weeks post-SE as shown in Fig. S1F-G, EZH2nHom mice received 10 doses of CP690550 from 7-9 weeks post-SE. **(B)** CP690550 reduced seizure frequency by 80% in EZH2nHom mice. EZH2nHom mice exhibited seizure frequencies 58% below WT mice (1-way ANOVA with Tukey’s correction; n=6-10). **(C)** Racine scale seizures were quantified as % total events (inset: cumulative frequency distribution). Daily CP690550 reduced seizure severity in nHom mice compared to pre-treatment (Wilcoxon matched pairs signed rank; W:-36.00, p<0.01, n=6) and WT mice (Mann Whitney test; U:119, p<0.05; n=6-10). **(D-E)** To assess the anticonvulsant properties of CP690550, naïve wildtype mice were pre-treated with CP690550 or vehicle (i.p., 15mg/kg) 30 minutes prior to threshold testing with flurothyl or PTZ (i.p., 45mg/kg). Time to generalized tonic clonic seizure was recorded. Results were analyzed by unpaired t-test. CP690550 pre-treatment did not alter **(F)** flurothyl (unpaired t-test; n=4-5) or **(G)** PTZ seizure threshold (unpaired t-test; n=5). *(**Seizure Model:** Systemic kainate; **Seizure detection:** Video recording)*

**Fig. S12.**
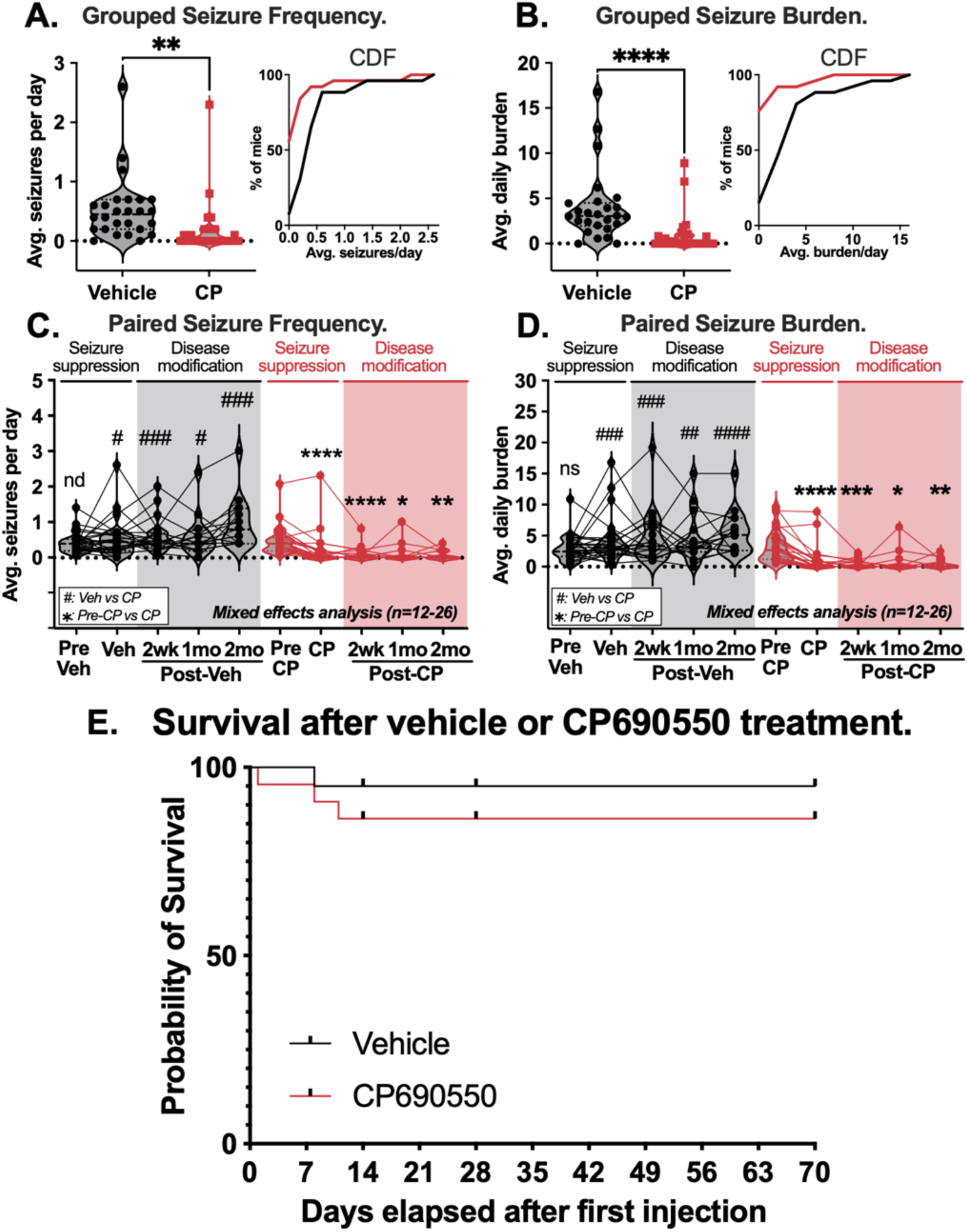
Long-term SRS data for both responders and non-responders to CP690550 in the systemic kainate model. SE was induced via systemic kainate, and behavioral seizures monitored by video from weeks 8-20 post-SE. Vehicle was administered from weeks 8-10 post-SE, followed by daily vehicle or CP690550 treatment weeks 11-12. After drug washout, video recording continued weeks 13-20 post-SE. **(A-B)** Seizure frequency and burden during treatment were displayed as violin plots and cumulative frequency distributions (bin centers are marked). Drug-treated mice showed reductions in seizure frequency and burden during CP690550 treatment (Kolmogorov-Smirnov test; **Frequency:** 3-fold, D:0.532, p<0.01, **Burden:** 4-fold, D:0.757, p<10^−4^; n=25-26). Paired seizure frequency **(C)** and burden **(D)** before treatment, during treatment, 2 weeks, 1 month, and 2 months post-washout were displayed as violin plots (✱: CP-treated compared to pre-CP, #: vehicle compared to CP-treated) (Mixed effects 2-way ANOVA with BKY correction; n=12-27). 2 weeks post-CP690550, mice showed a 6-fold reduction in frequency (q<10^−3^) and a 11-fold reduction in burden (q<10^−3^) compared to 2 weeks post-vehicle; 12 out of 18 mice (67%) were seizure-free during this time. At 2 months post-washout, drug-treated animals continued to exhibit a 12-fold reduction in seizure frequency (q<0.01) and an 11-fold reduction in seizure burden (q<0.01) compared to vehicle-treated mice (n=12-15). **(E)** Of the 55 male and female FVB mice treated with vehicle or CP690550, 51 survived treatment. One vehicle-treated mouse died 8 days after the the first day of treatment. Three CP690550-treated mice died 1, 8, and 11 days after the first day of treatment. 13 mice were harvested on the final day of treatment (n=6 vehicle, 7 CP690550). 11 mice were harvested two weeks post-washout (n=5 vehicle, 6 CP690550). 12 mice were harvested two weeks post-washout (n=7 vehicle, 5 CP690550). There was no difference in survival between treatment groups (Log-rank Mantel-Cox test; Chi square: 0.893, p=0.344). *(**Seizure Model:** Systemic kainate; **Seizure detection:** Video recording)*

**Fig. S13.**
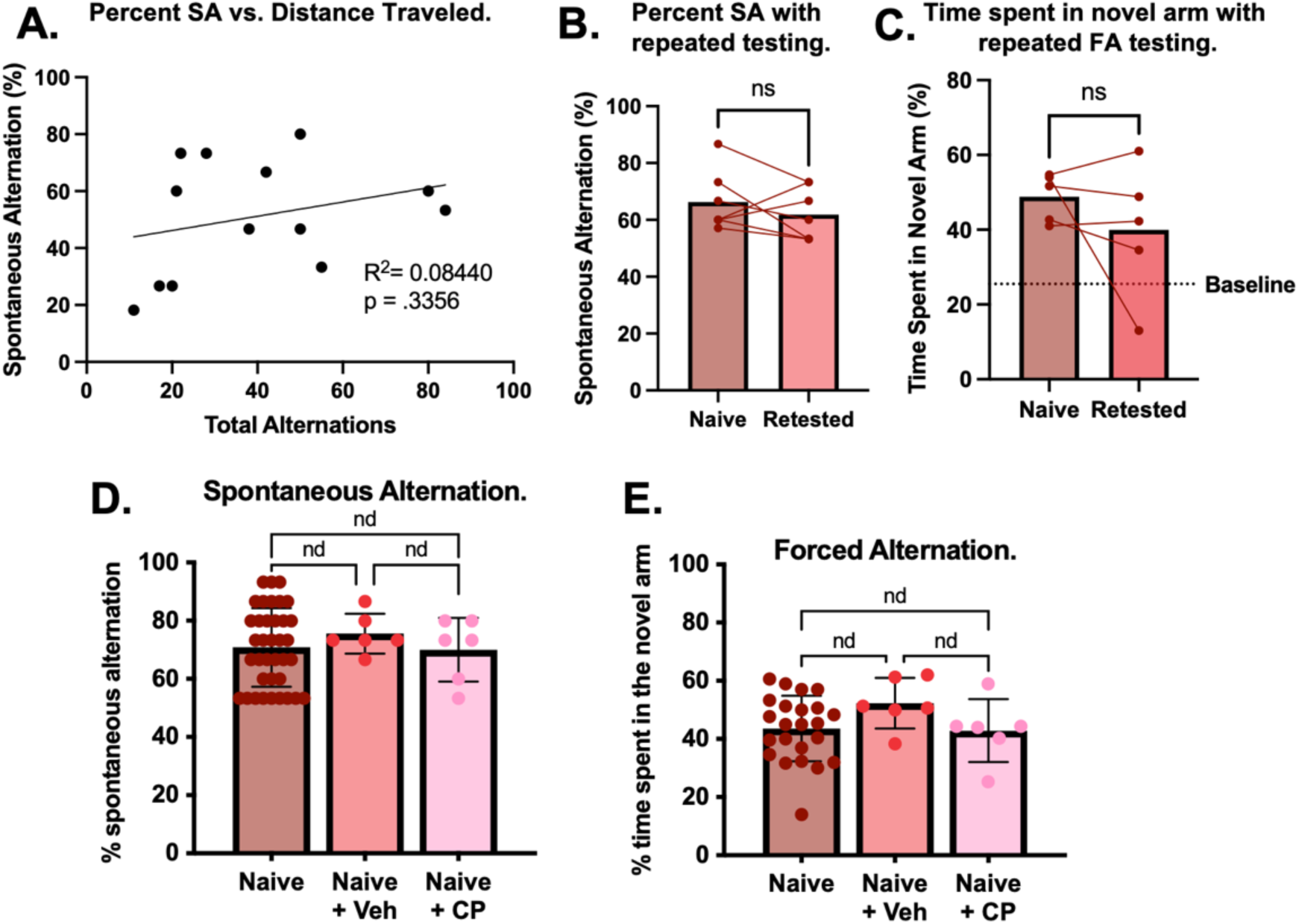
Distance traveled is not correlated with deficits in spontaneous alternation post-SE, and SA and FA results are not altered with repeated testing or CP690550 treatment in naïve mice. **(A)** Epilepsy-related reductions in working memory (as measured by percent spontaneous alternations) were not correlated to distance traveled as measured by the total number of alternations (Pearson correlation; n=13). Percent spontaneous alternation **(B)** and time spent in the novel arm during the retrieval trial of FA testing **(C)** were not altered by repeated testing (paired t-test; n=5-6). The line demarcating baseline in panel C was found by measuring the amount of time mice spent in a random arm of the maze without ever having been exposed to the sample trial (i.e., the trial in which one of the arms was blocked). Baseline was ∼25% instead of the theoretically expected 33% based on the criteria by which a mouse was considered to be “in” versus “out” of an arm; time in the center of the maze was not counted (See Methods). Tests occurred 24 days apart for spontaneous alternation, and 17 days apart for forced alternation. SA **(D)** and FA **(E)** tests were performed on untreated naïve mice, and mice treated with vehicle or CP690550 for two weeks (i.p., 15mg/kg/day). Neither vehicle nor CP treatment altered percent spontaneous alternation or time spent in the novel arm. Panels D-E were analyzed by 1-way ANOVA with BKY’s correction for multiple comparisons (**SA:** n=6-37; **FA:** n=6-23). *(**Seizure Model:** Systemic kainate)*

**Fig. S14.**
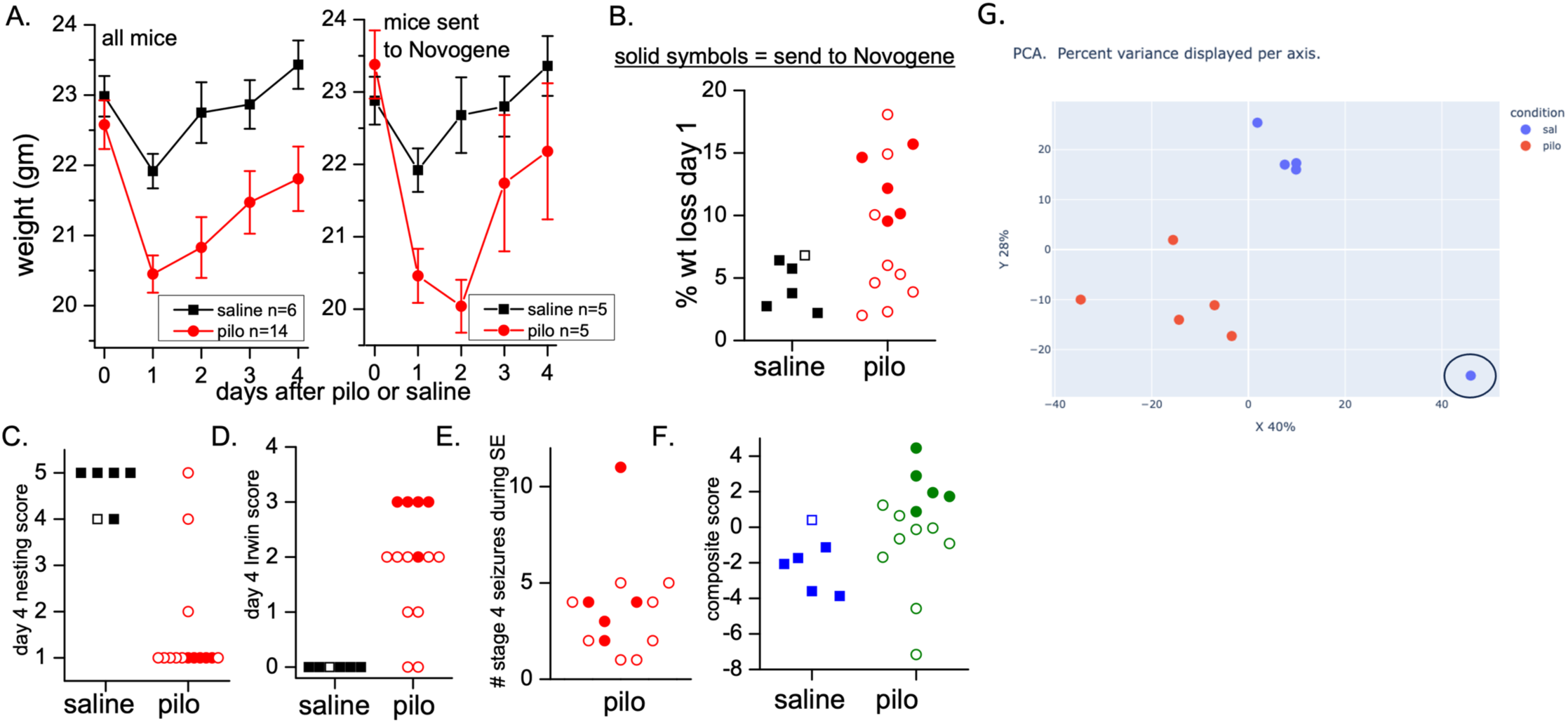
Selection of mice for snRNAseq and Principal Component Analysis on HVGs. **(A)** Weights of saline- and pilocarpine-injected mice were tracked up to four days post-SE for (left) all mice and (right) mice sent to Novogene. **(B)** Percent weight loss on day 1 post-SE in saline versus pilo mice; solid symbols denote mice sent to Novogene from each treatment group. On day 4 post-SE, **(C)** nesting scores and **(D)** Irwin scores for neurobehavioral recovery were recorded. **(E)** The number of Racine class IV seizures during pilocarpine injection were tabulated for each animal. **(F)** The mean number of standard deviations from the mean of each measure was calculated across all 6 saline-treated and 14 pilocarpine-treated mice to generate a composite score for each animal, and mice were selected based on wide separation between saline and pilo groups. **(G)** The 4000 most variant genes were selected to subject samples to PCA analysis. Four saline samples clustered together away from the pilo samples. One saline sample did not cluster with either group and was discarded (circled blue dot). This resulted in 5 pilo samples, 4 saline samples, and 58,365 cells. *(**Seizure Model:** Pilocarpine)*

## Supplemental Tables

**Table S1.** Tabulated daily seizure frequency, burden, and metadata for responders and non-responders to CP690550 in the systemic kainate model.

**Table S2.** Fisher Exact test results for mouse vs. human cluster overlap analysis.

**Table S3.** Mouse strain, seizure model, and seizure detection method information for each experiment.

